# Age, sex, and vendor contributions to variance in Diffusion Tensor Imaging (DTI) ‘Big Data’

**DOI:** 10.64898/2026.04.28.721286

**Authors:** Nicholas Simard, Michael D Noseworthy

## Abstract

The aim of this study was to evaluate the contributions of age, sex, and MRI vendor to variance in Diffusion Tensor Imaging (DTI) metrics, with a focus on understanding the impact of these factors in large-scale healthy brain datasets. A dataset of 2,700 DTI scans from healthy controls across multiple sites and MRI vendors was analyzed. The DTI scalar metrics fractional anisotropy (FA) and mean diffusivity (MD) were processed and the influence of age, sex, vendor, and brain atlas selection were determined. A statistical analysis was conducted and revealed significant (p<0.05) age-related differences in DTI metrics, with older participants showing reduced FA and increased MD, in line with known microstructural changes. Sex differences were observed, with females exhibiting slightly higher FA and lower MD in certain brain regions. Vendor variability was also noted, with all three MRI vendors showing significant differences in FA with Siemens machines typically exhibiting higher FA values and GE machines lower FA values (i.e. FA_Siemens_ > FA_Philips_ > FA_GE_). Atlas selection also highlighted some specific ROI behaviour (e.g. tapetum of the corpus callosum) as one of the most significant regions of interest (ROIs) in the JHU-Tracts atlas that demonstrated a large amount of deterioration with age, particularly in females. These findings emphasize the need to account for biological factors such as age and sex, as well as technical factors like ROI selection and MRI vendor, when interpreting DTI data. The results demonstrate the potential of large-scale, multi-vendor datasets to uncover meaningful biological trends, while also addressing the challenges of scanner-specific variability. Although previous work has shown sex and age differences, this is the first large scale DTI analysis that has included age, sex, and MRI vendor as sources of variance in one model.

## Introduction

Diffusion Tensor Imaging (DTI) is a valuable neuroimaging technique that provides non-invasive insight into the structural integrity of white matter **[1]**. Widely used in clinical and research contexts, DTI has applications ranging from preoperative neurosurgical planning **[2,3]** to understanding neurodegenerative diseases **[4]**. Quantitative DTI metrics, such as fractional anisotropy (FA) and mean diffusivity (MD), are increasingly recognized as valuable biomarkers for brain health and disease **[5,6]**. However, the reliability and interpretability of these metrics are influenced by several factors, including biological variables such as age **[7,8,9]** and sex **[8,10,11]**, as well as technical differences across MRI vendors **[12,13,14]**) and regions of interest (ROIs) **[12,15]**.

Biological effects of aging were first characterized through imaging modalities by evaluating cerebrospinal fluid (CSF), gray matter (GM), and white matter (WM) volume changes in the brain **[16,17,18]**. Age-related changes in WM microstructure have since been validated in the context of DTI with studies reporting decreases in FA and increases in MD as hallmarks of aging **[7,8,9,19,20]**. While conventional DTI research has primarily focused on increases in FA due to white matter maturation in adolescence **[21,22]**, the processes underlying healthy aging remains underexplored and warrants greater attention to understand its complex nature **[21,23,24]**. Similarly, sex differences in WM organization have been identified in specific brain regions such as the corpus callosum (CC), where males tend to exhibit higher FA values compared to females **[8,10,11]**, whereas females exhibit higher MD values in the superior longitudinal fasciculus (SLF) **[25]**. Variability in biological measures are often further increased by differences in MRI scanner vendors where phantom studies show limited differences (∼1.7-2%) and high reliability across platforms **[25,26,27]**. Yet, human studies demonstrate that Philips scanners exhibit lower FA values **[13]** compared to traditionally higher FA values observed in Siemens scanners **[14,24]**. There are also technical nuances and considerations that can induce variability in DTI metrics, particularly the selection of ROIs **[15]**. There can be variability due to manually selecting ROIs **[28]**, using atlas-based ROIs with thresholding **[29,30]**, or even when opting for voxel-based methods **[28]**. Different MRI manufacturers can also cause DTI measurements to change **[12]**. This variability is due to a variety of reasons such as different proprietary motion probing gradients (MPGs), warping effects due to registration, and partial voluming effects **[28]**. Addressing and understanding these sources of variance is critical for leveraging large-scale DTI datasets to uncover meaningful trends.

The advent of multi-site neuroimaging repositories has further enabled the study of age, sex, vendor, and ROI variability across diverse populations **[31,32,33,34,35,36]**. The increasing availability of DTI data, coupled with large sample sizes, provides the statistical power necessary to investigate these variability effects on a practical scale **[37,38]**. DTI’s increasing richness has made it a valuable resource in machine learning (ML) and artificial intelligence (AI) applications, where large-scale datasets are essential for training robust algorithms **[39,40]**.

This study aimed to quantify the contributions of variance in DTI data due to *Age*, *Sex*, *Vendor*, and *Atlas* in the context of pre-selected ROIs within a large heterogeneous dataset. Sourcing 2,700 DTI healthy control datasets scanned across multiple sites and found within multiple repositories, we applied robust statistical methods to evaluate the relative impact of these factors on FA and MD. By disentangling biological effects from technical variability, this work provides critical insights into the understanding of DTI scalar metric changes and highlights the nuances required to appropriately adopt large-scale datasets for ML/AI applications and for advancing our understanding of brain microstructure **[39,40,41,42,43]**.

## Material & Methods

### Dataset Description

This study utilized a curated sample of 2,700 diffusion tensor imaging (DTI) datasets of healthy controls that were downloaded from several global open-source repositories ensuring substantial heterogeneity of the data sample along with proper diversity representation **[44]**. Data was downloaded from the Human Connectome Project (HCP) (https://db.humanconnectome.org) **[32]**, the Adolescent Brain Cognitive Development (ABCD) study (https://abcdstudy.org/) **[45]**, the Imaging Data Archive (IDA), managed by the Laboratory Neuro Imaging (LONI) (https://ida.loni.usc.edu/) **[33]**, the Neuroimaging Tools and Resources Collaboratory (NITRC) (https://www.nitrc.org) **[9]**, the enhanced Nathan Kline Institute-Rockland Sample (NKI-RS) (https://www.nki.rfmh.org) **[38]**, the OASIS-3 Longitudinal Neuroimaging, Clinical, and Cognitive Dataset for Normal Aging and Alzheimer’s Disease (https://sites.wustl.edu/oasisbrains/home/oasis-3/) **[46]**, and the OpenNeuro resource for sharing of neuroscience data (https://openneuro.org) **[47]**. The datasets sourced included balanced males and females between the ages of 20-65 with *Age* being further segmented into 5-year bins starting at age 20-25 and ending in 61-65 (9 bins in total). Data from all three major *Vendors* (GE, Siemens, Philips) were collected exclusively on 3T machines. The resulting data distribution was equivalent to 50 datasets for each *Age*, *Vendor*, and *Sex* (i.e. 50×3×9×2), resulting in a total of 2,700 samples. Subjects were selected as healthy controls, meaning no previous brain disease, no known neurological/neurocognitive disorder, no previous head injury, or no DSM-IV psychiatric diagnosis **[48,49]**. This study was approved by our local research ethics board (Hamilton Integrated Research Ethics Board, HiREB. Project #18683).

### Image Acquisition Parameters

All datasets were acquired using 3T MRI systems across multiple sites, representing a broad range of scanning environments. Also, as using more gradient directions improves the precision and accuracy of the FA measurement, only datasets with 30 or more gradient directions were sourced for the study **[50]**. The MRI systems investigated in this study included the big three vendors (GE, Siemens, and Philips), ensuring diversity in scanner type and technological capabilities. For each data acquisition, different receiver coil arrays were employed, including 8-channel, 16-channel, or 32-channel brain phased array coils. A variety of scanning parameters were utilized across the different systems including variations in echo time (TE), repetition time (TR), gradient directions, and number/strength of b-values, among others (**Table 1**). These differing parameters provide a comprehensive overview of the technical specifications used in this study, highlighting the variability and inhomogeneity in scanning protocols across the datasets.

**Table 1.**
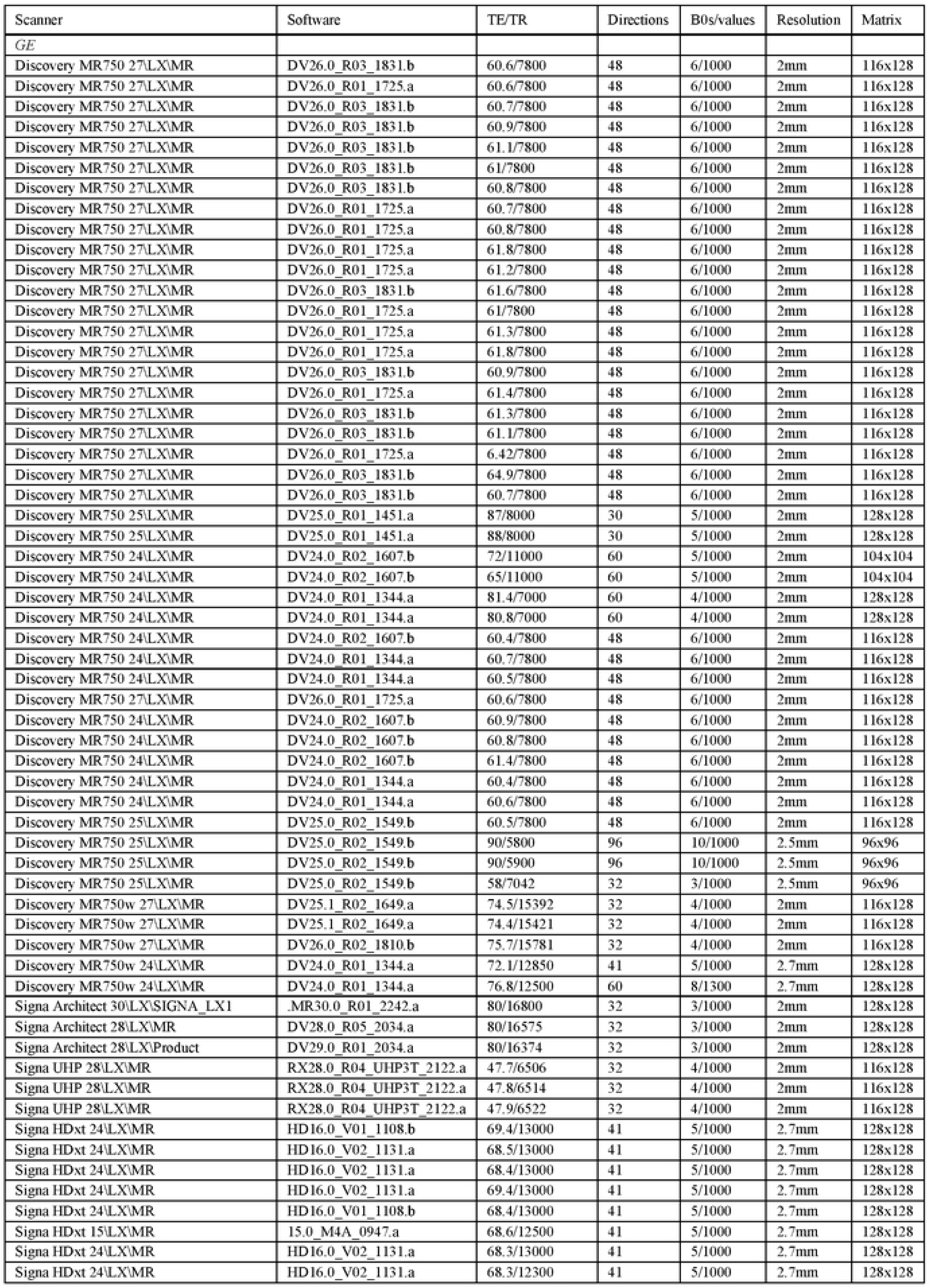
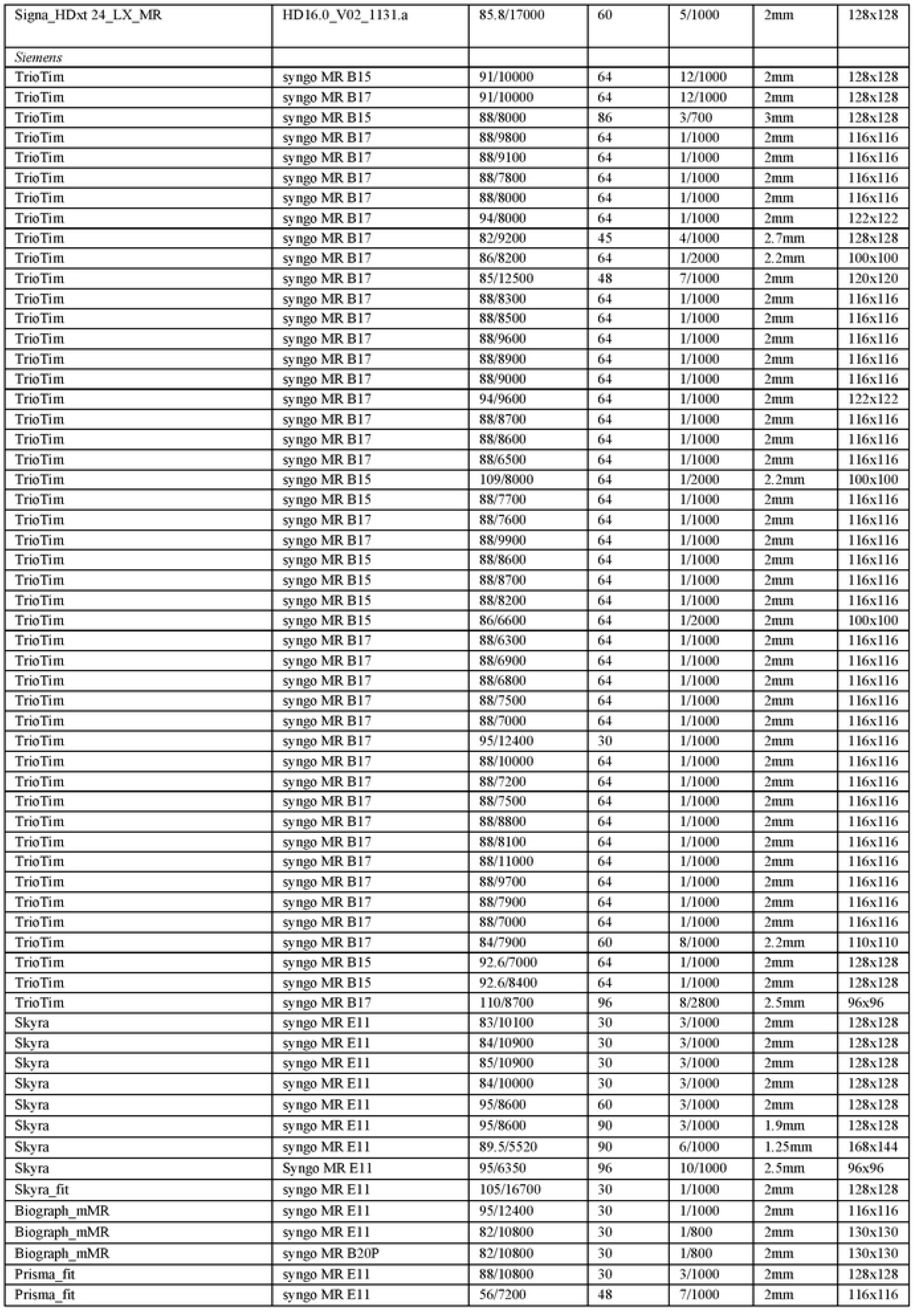
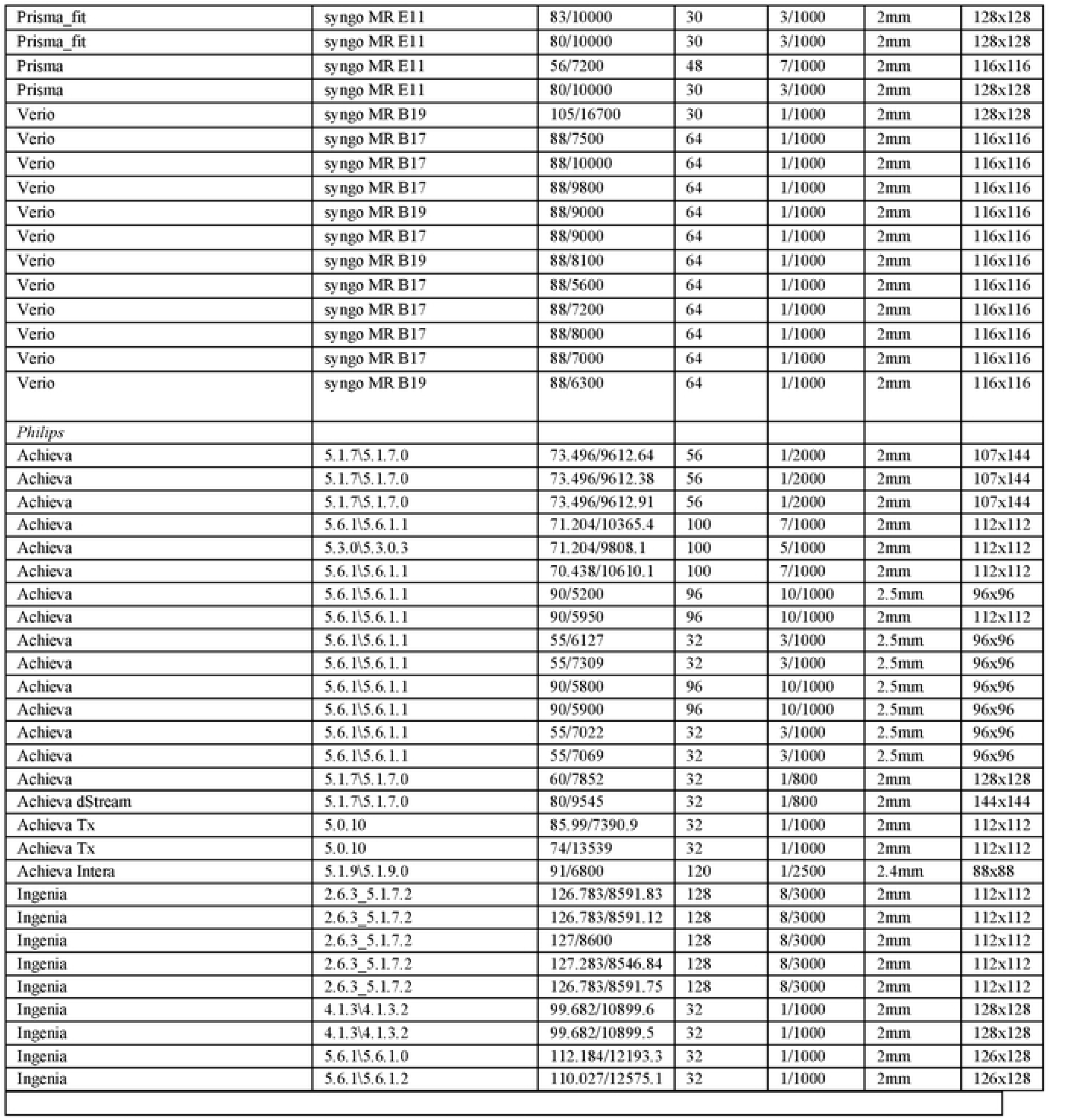
This table displays the technical specifications of MRI scanners across different vendors and platforms. Detailed characteristics of each MRI scanner used in this study including scanner type, software version, TE/TR values, number of diffusion directions, b-values, resolution, and matrix. The specifications cover big name manufacturers GE, Siemens, and Philips, highlighting the diversity and variability in scanning configurations from open-access repositories. The goal is to outline the diversity and inhomogeneity and understand the implications these technical differences may have on MRI data consistency and reproducibility.

### Image Processing

DTI data was processed using a standardized pipeline integrating *fsl* (FMRIB Software Library, version 5.0.11) tools for robust pre-processing and metric extraction **[51]**. Initial steps included brain extraction using *bet2* **[52]** and eddy current distortion correction with *eddy_correct* **[53]** to minimize artifacts. All datasets were co-registered to the MNI152 T1-weighted 1mm standard brain space using *flirt* **[54,55,56]** with a 12 degree-of-freedom affine registration, ensuring spatial consistency across subjects and imaging vendors. For enhanced accuracy in warping, nonlinear registration was performed using the *fnirt* tool, employing a b-spline representation to refine the transformation into the standard space. DTI metric maps, including fractional anisotropy (FA), mean diffusivity (MD), radial diffusivity (RD), and axial diffusivity (AD), were generated using *dtifit* **[57,58]**. FA, as a primary metric, was prioritized due to its demonstrated sensitivity in detecting WM integrity alterations. The diffusivity maps provided a comprehensive assessment of structural variability and facilitated robust comparisons across subjects, accounting for both biological variability (*Age* & *Sex*) and technical differences introduced by scanner variations.

### Regions of Interest (ROIs)

*Age*, *Sex*, and *Vendor* factors all affect the brain’s microstructure in different ways and to a different degree, therefore regional differences were examined in detail as a crucial part of this study. For the DTI metric maps (FA, MD), ROIs were carefully selected to provide full WM coverage. As a rank-2 tensor was calculated, ROIs pertaining to predetermined tracts help ensure that predominant fiber tract directions exist and helps minimize partial voluming effects **[30]**. However, it is often noted that some ROIs still demonstrate high amounts of variability due to *Age* and *Sex*, such as the uncinate fasciculus (UF) **[59]** and the CC **[21,22,60]**. For this reason, three distinct *Atlases* were selected to evaluate the variability in this study including the JHU-Labels, JHU-Tracts, and XTRACT atlases (**Table 2**). The JHU-Labels *Atlas* is a subset of 50 WM tracts from the JHU ICBM-DTI-81 atlas that provides anatomically defined ROIs widely validated and frequently used in DTI studies to examine WM integrity across standard regions **[61,62,63]**. The JHU-Tracts atlas is another subset of 20 WM tracts from the JHU ICBM-DTI-81 atlas that complements the JHU-Labels atlas with representations of larger fiber tracts **[61,62,63]**. The XTRACT atlas was included for its comprehensive set of 42 expertly delineated fiber pathways, extending coverage to tracts that are critical for functional and structural connectivity analyses but are less commonly examined in traditional atlases **[64]**. All *Atlases* were subject to 90\% binarized thresholding to ensure appropriate ROI coverage before proceeding with analysis. The multi-atlas approach was selected to mitigate the effects of partial voluming and ensure high anatomical fidelity in regional analyses.

**Table 2.**
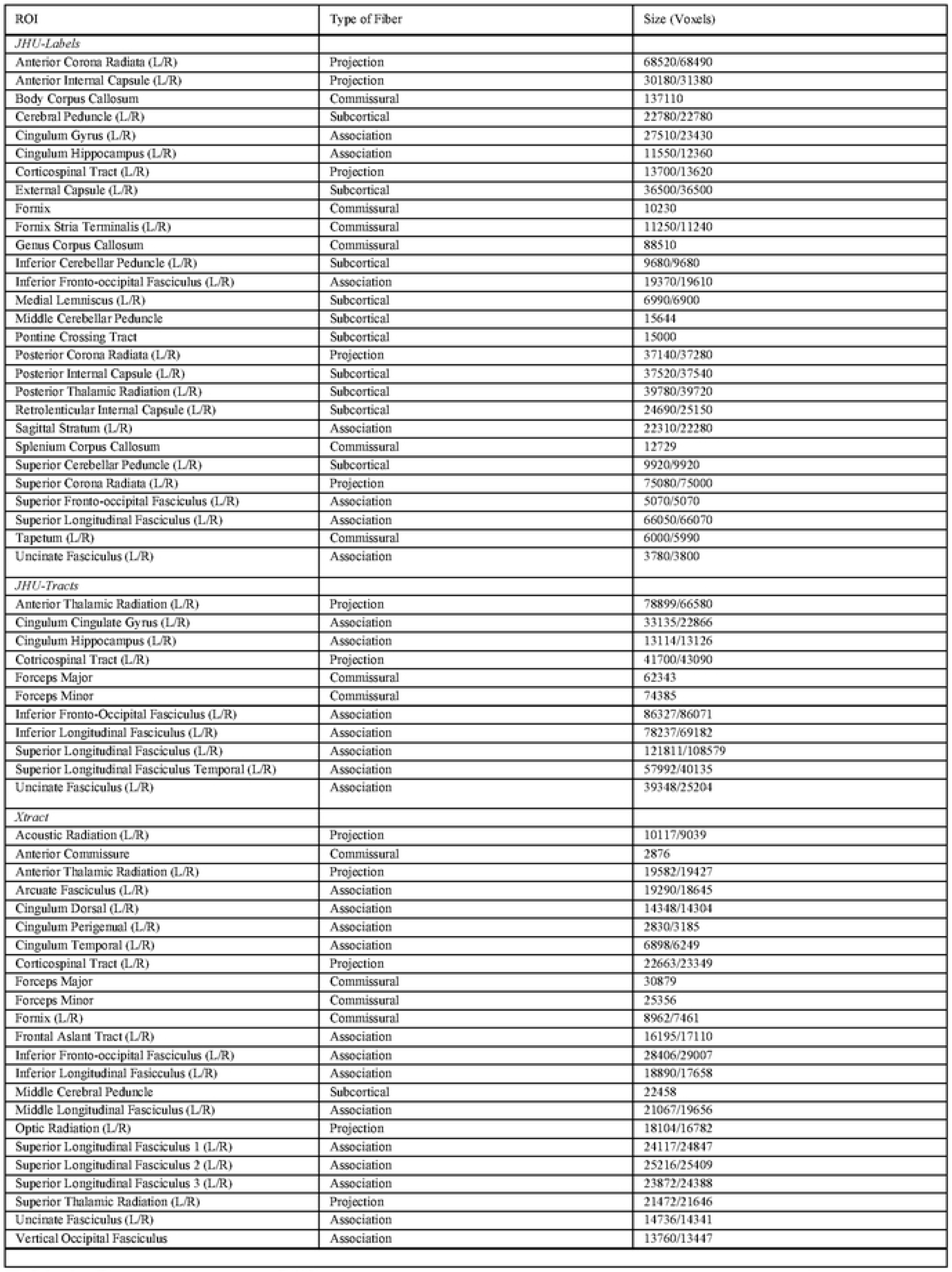
Region of interest *(ROI) Atlas* selections with corresponding fiber type classifications and voxel volumes. *ROIs* were selected from three traditional atlases including the JHU-Labels, JHU-Tracts, and XTRACT *Atlases* which are utilized in neuroanatomical analysis. The specific fiber types are categorized as projection, commissural, or association fibers. The size of each *ROI* is indicated in voxel volume (1mm’ voxel units) and shown for both left (L) and right (R) hemispheres, where applicable.

### Statistical Analysis

Once all the data was preprocessed, the statistical analysis focused on 4 factors evaluating the variability of *Age*, *Sex*, *ROI*, and *Vendor*. Each discrete *Age* factor was analyzed (20-65) **[20]**.

However, age group binning (20-25, 26-30, 31-35, 35-40, 41-45, 46-50, 51-55, 56-60, 61-65) was used to evaluate the variability effects of *Age*, including the global percentage change per *Age* bin **[65]**. For *Sex*, a binarized male vs. female investigation was performed. For *ROI*, each of the 112 *ROIs*, along with the three *Atlases* themselves were investigated for variability. Principal component analysis (PCA) was also performed to identify specific *ROIs* that contribute to the most variance. *ROI* was also a target variable to understand how the interactions with *Age*, *Sex*, and *Vendor* influence specific *ROI* measurements. For *Vendor*, an investigation on factors GE, Siemens, and Philips was performed. For all factors, heatmaps, 4-way ANOVA, and coefficient of variation (CV) **[59]** analysis was performed to better understand the impact of variability due to each factor. A series of interaction and circular bar plots were also generated to demonstrate the effects of each factor within the context of FA, whereas a series of bar plots demonstrated the effects of each factor on AD, MD, and RD. Furthermore, the *3dANOVA* command in AFNI, generated with 4 levels (or factors) demonstrated the areas of high variability in WM, independent of *ROI* analysis **[66]**.

## Results

### Global Effects on White Matter Variability

The global variability of WM within the context of ‘Big Data’ was significantly affected by the investigated factors of *Age*, *Sex*, *Vendor*, and *Atlas* within the context of *ROI*. Literature has shown that WM integrity decreases with aging (post-puberty), exhibiting lower FA values and higher MD, AD, and RD values **[20,65]**, particularly at the fifth decade **[67]**. In this study (**Fig. 1**), *ROIs* were pooled rather than treated as a separate factor and *Age* was investigated as a continuous variable showing global WM integrity following typical behaviour with significant *Age*-related decreases in FA (r = 0.755, p < 0.0001) compared to significant increases in AD (r = 0.545, p < 0.001), MD (r = 0.582, p < 0.001), and RD (r = 0.615, p < 0.001). Investigating *Age* through 5-year binned groups (**Fig. 1**), with 20-25 year olds as the reference, also showcased a global FA percentage change decrease due to Age (µ_Female_ =2.696, µ_Male_ =4.108) compared to the global AD (µ_Female_ = 1.800, µ_Male_ = 2.072), MD (µ_Female_ = 3.00, µ_Male_ = 2.727), and RD (µ_Female_ = 4.210, µ_Male_ = 3.496) increases. For some cases in the 26-30 and 31-35 age groups, there are increases in FA or decreases in AD, MD, RD, which is likely explained by the variability of puberty into late adolescence with some degree of WM maturation **[21,22]**. The global FA percentage change also starts to show *Sex* differences in which females are more resilient to *Age*-related changes compared to males. A 4-way ANOVA with *Age* (5-year bins), *Sex*, *Vendor*, and *ROI* as factors (**Table 3**), also demonstrates the significant level of variability associated with *Age* (all p < 0.001) throughout all *Atlases* and *ROIs*. Lastly, investigating the *Age* (5-year bins) factor as a function of FA maps (**Fig. 2**) demonstrates the areas of significant variability (p < 0.001). High levels of variability are found in the interhemispheric communication regions such as the CC, matching the literature **[65]**.

**Figure 1.**
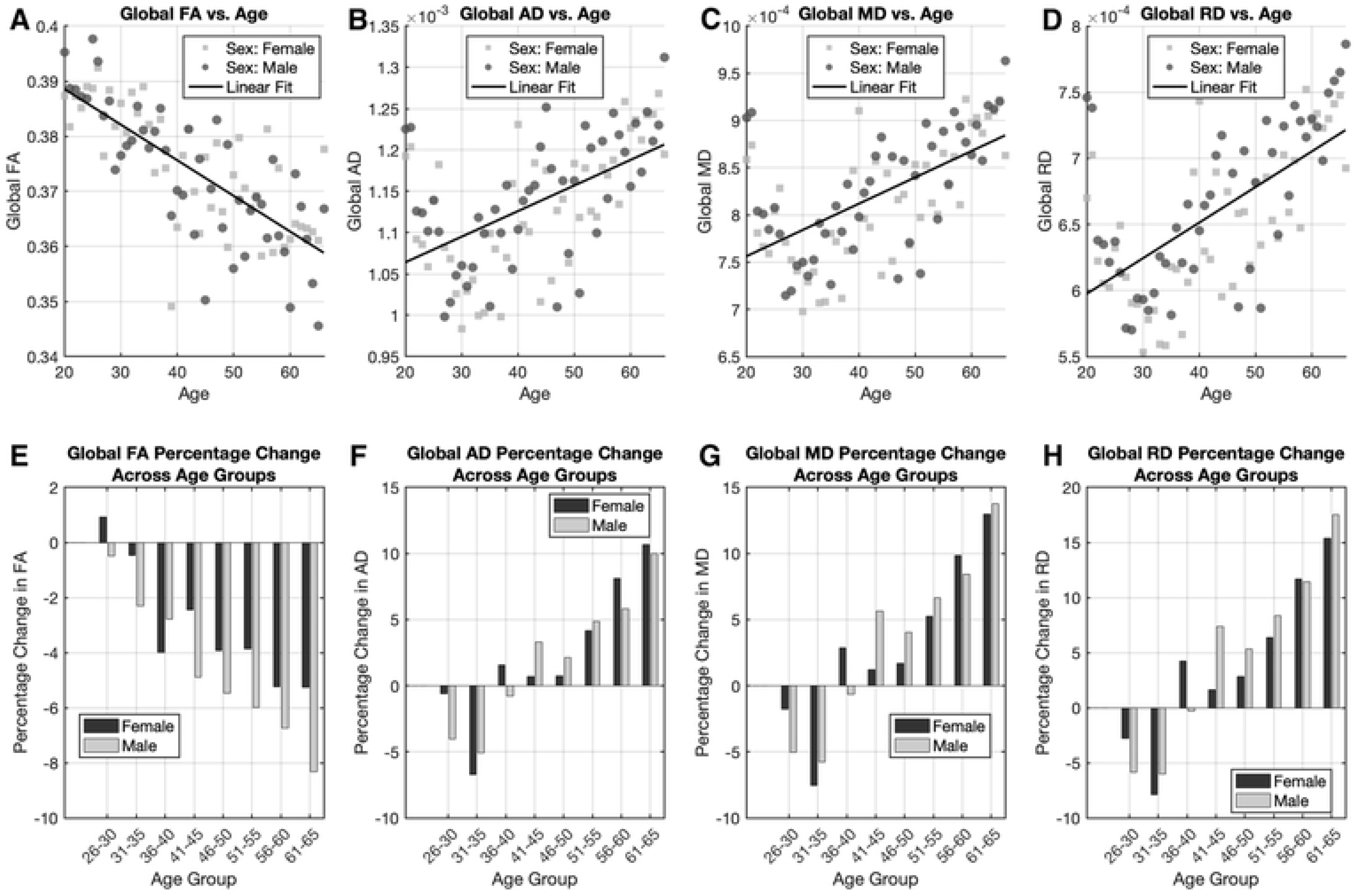
Scatterplots pooling measures from all *ROIs* to depict global diffusion measures as a function of *Age* across male and female participants: (A) Fractional Anisotropy (FA), (B) Axial Diffusivity (AD), (C) Mean Diffusivity (MD), and (D) Radial Diffusivity (RD). Male data points are identified in dark gray filled circles while female data points are identified in light gray squares. Trend lines highlight *Age*-related changes in each measure with *Age* as a continuous variable. (E-H) Bar graphs illustrating the percentage changes in diffusion measures across all *ROIs* per 5-year binned *Age* using 20-25 year olds as a reference: (E) FA, (F) AD, (G) MD, and (H) RD. These plots summarize the relative differences in diffusion metrics between *Age*, providing insights into their nonlinear progression across the lifespan. Male and female data points are also shown in these bar graphs to highlight *Sex* differences related to *Age*.

**Figure 2.**
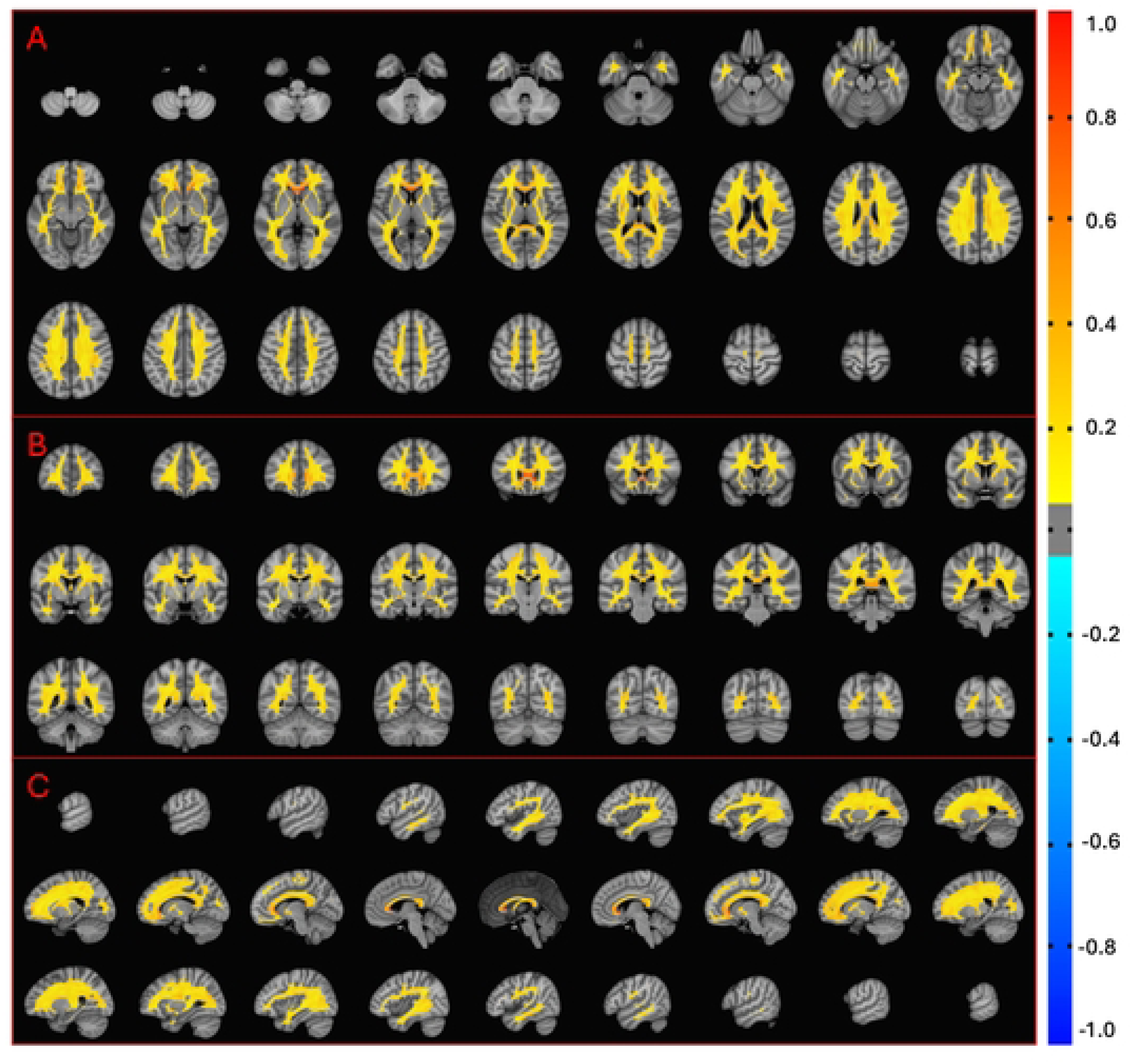
Variability in Fractional Anisotropy (FA) across the brain due to *Age*, visualized using a 9 × 3 grid of axial (A), coronal (B), and sagittal (C) slices from a *3dANOVA* analysis in AFNI **[66]**. Analysis was performed on FA maps of all individuals (𝑛 = 2700) stratified in their respective 5-year *Age* bins. Regions with higher FA variability are shown in orange, particularly concentrated near interhemispheric areas such as the corpus callosum (CC), indicating pronounced *Age*-related changes in these areas. Yellow regions represent significant but less extreme variability, reflecting more moderate *Age*-related alterations. This visualization highlights the heterogeneous effects of *Age* on white matter (WM) integrity across the brain.

**Table 3.** Results of statistical analysis for *age, Sex,* Region of interest *(ROI),* and *Vendor* as factors across multiple *Aliases* showing the variability associated with diffusion tensor imaging (DTI) metrics. The table summarizes the F-critical (F), p-values, and effect sizes (η^2^) for the analysis of fractional anisotropy (FA), mean diffusivity (MD), axial diffusivity (AD), and radial diffusivity (RD) across different *Atlases-.* JHU-Labels, JHU-Tracts, XTRACT, and all *Atlases.* Significant measures are in bold where all F-critical values were significant along with all p-values (< 0.001), whereas effect sizes (η^2^), indicating the proportion of variance explained, varied in significance (η^2^ > 0.06). F-critical values of significance varied for JHU-Labels (1.121), JHU-Tracts (1.196), XTRACT (1.132), and all *Aliases* (1.080).

|  | JHU-Labels |  |  |  | JHU-Tracts |  |  |  | XTRACT |  |  |  | All Atlases |  |  |  |
| --- | --- | --- | --- | --- | --- | --- | --- | --- | --- | --- | --- | --- | --- | --- | --- | --- |
|  | *FA* | *MD* | *AD* | *RD* | *FA* | *MD* | *AD* | *RD* | *FA* | *MD* | *AD* | *RD* | *FA* | *MD* | *AD* | *RD* |
| **Age** |  |  |  |  |  |  |  |  |  |  |  |  |  |  |  |  |
| F | 514.6 | 894.2 | 840.4 | 884.7 | 530.1 | 846.4 | 685.4 | 958.2 | 784.7 | 1353 | 1096 | 1483 | 1367 | 2495 | 2222 | 2562 |
| P-value | .0001 | .0001 | .0001 | .0001 | .0001 | .0001 | .0001 | .0001 | .0001 | .0001 | .0001 | .0001 | .0001 | .0001 | .0001 | .0001 |
| $\eta^2$ | .0149 | .0282 | .0237 | .0293 | .0258 | .0833 | .0651 | .0920 | .0169 | .0578 | .0455 | .0588 | .0096 | .0367 | .0289 | .0383 |
| **Sex** |  |  |  |  |  |  |  |  |  |  |  |  |  |  |  |  |
| F | 179 | 362.9 | 159.0 | 503.2 | 266.0 | 55.39 | 31.28 | 75.05 | 55.38 | 152.9 | 67.32 | 216.6 | 23.57 | 566.7 | 255.4 | 785.2 |
| P-value | .0001 | .0001 | .0001 | .0001 | .0001 | .0001 | .0001 | .0001 | .0001 | .0001 | .0001 | .0001 | .0001 | .0001 | .0001 | .0001 |
| $\eta^2$ | .0006 | .0015 | .0005 | .0018 | .0016 | .0001 | .0001 | .0001 | .0001 | .0001 | .0001 | .0001 | .0001 | .0011 | .0004 | .0013 |
| **ROI** |  |  |  |  |  |  |  |  |  |  |  |  |  |  |  |  |
| F | 2509 | 1829 | 2296 | 1586 | 5169 | 306.1 | 451.7 | 305.9 | 5853 | 979.1 | 937.5 | 1213 | 7159 | 1516 | 2049 | 1471 |
| P-value | .0001 | .0001 | .0001 | .0001 | .0001 | .0001 | .0001 | .0001 | .0001 | .0001 | .0001 | .0001 | .0001 | .0001 | .0001 | .0001 |
| $\eta^2$ | .4437 | .3531 | .3996 | .3256 | .5985 | .0741 | .1065 | .0690 | .6442 | .2110 | .2036 | .2516 | .6959 | .3079 | .3703 | .3056 |
| **Vendor** |  |  |  |  |  |  |  |  |  |  |  |  |  |  |  |  |
| F | 1174 | 2946 | 4404 | 2167 | 1392 | 2570 | 2783 | 2446 | 1439 | 3840 | 4409 | 3370 | 2867 | 7763 | 9031 | 6190 |
| P-value | .0001 | .0001 | .0001 | .0001 | .0001 | .0001 | .0001 | .0001 | .0001 | .0001 | .0001 | .0001 | .0001 | .0001 | .0001 | .0001 |
| $\eta^2$ | .0085 | .0228 | .0314 | .0177 | .0170 | .0648 | .0651 | .0575 | .0072 | .0405 | .0474 | .0327 | .0050 | .0282 | .0337 | .0234 |
| **Age-Sex** |  |  |  |  |  |  |  |  |  |  |  |  |  |  |  |  |
| F | 50.76 | 51.05 | 41.94 | 52.38 | 31.11 | 30.97 | 26.47 | 34.01 | 37.82 | 53.16 | 47.91 | 55.31 | 91.46 | 114.5 | 99.92 | 116.1 |
| P-value | .0001 | .0001 | .0001 | .0001 | .0001 | .0001 | .0001 | .0001 | .0001 | .0001 | .0001 | .0001 | .0001 | .0001 | .0001 | .0001 |
| $\eta^2$ | .0015 | .0015 | .0010 | .0018 | .0015 | .0001 | .0001 | .0001 | .0008 | .0029 | .0020 | .0033 | .0006 | .0017 | .0015 | .0019 |
| **Age-ROI** |  |  |  |  |  |  |  |  |  |  |  |  |  |  |  |  |
| F | 13.76 | 14.92 | 12.03 | 16.45 | 9.270 | 1.930 | 1.550 | 2.690 | 10.20 | 3.910 | 3.150 | 4.750 | 13.12 | 11.92 | 9.520 | 13.39 |
| P-value | .0001 | .0001 | .0001 | .0001 | .0001 | .0001 | .0001 | .0001 | .0001 | .0001 | .0001 | .0001 | .0001 | .0001 | .0001 | .0001 |
| $\eta^2$ | .0195 | .0228 | .0170 | .0266 | .0086 | .0001 | .0001 | .0001 | .0090 | .0058 | .0059 | .0065 | .0102 | .0192 | .0139 | .0221 |
| **Age-Vendor** |  |  |  |  |  |  |  |  |  |  |  |  |  |  |  |  |
| F | 73.34 | 399.5 | 573.4 | 269.1 | 80.77 | 443.6 | 495.9 | 384.9 | 126.7 | 827.3 | 1002 | 655.0 | 183.8 | 1272 | 1698 | 920.7 |
| P-value | .0001 | .0001 | .0001 | .0001 | .0001 | .0001 | .0001 | .0001 | .0001 | .0001 | .0001 | .0001 | .0001 | .0001 | .0001 | .0001 |
| $\eta^2$ | .0042 | .0251 | .0324 | .0177 | .0079 | .0926 | .0947 | .0805 | .0054 | .0694 | .0850 | .0523 | .0026 | .0373 | .0443 | .0279 |
| **Sex-ROI** |  |  |  |  |  |  |  |  |  |  |  |  |  |  |  |  |
| F | 30.60 | 16.52 | 11.36 | 20.60 | 6.540 | 2.290 | 1.66 | 3.000 | 8.560 | 5.790 | 4.230 | 6.670 | 28.03 | 14.11 | 9.250 | 18.02 |
| P-value | .0001 | .0001 | .0001 | .0001 | .0001 | .0011 | .0347 | .0011 | .0001 | .0001 | .0001 | .0001 | .0001 | .0001 | .0001 | .0001 |
| $\eta^2$ | .0054 | .0030 | .0021 | .0044 | .0008 | .0001 | .0001 | .0001 | .0009 | .0001 | .0001 | .0001 | .0027 | .0028 | .0018 | .0039 |
| **Sex-Vendor** |  |  |  |  |  |  |  |  |  |  |  |  |  |  |  |  |
| F | 36.80 | 60.76 | 64.34 | 73.95 | 111.0 | 85.45 | 94.24 | 93.26 | 53.72 | 135.4 | 132.2 | 153.2 | 91.02 | 207.5 | 216.3 | 236.8 |
| P-value | .0001 | .0001 | .0001 | .0001 | .0001 | .0001 | .0001 | .0001 | .0001 | .0001 | .0001 | .0001 | .0001 | .0001 | .0001 | .0001 |
| $\eta^2$ | .0003 | .0008 | .0005 | 0.0009 | .0014 | .0001 | .0001 | .0001 | .0003 | .0001 | .0020 | .0001 | .0002 | .0006 | .0007 | .0006 |
| **ROI-Vendor** |  |  |  |  |  |  |  |  |  |  |  |  |  |  |  |  |
| F | 46.72 | 60.76 | 42.64 | 26.75 | 40.88 | 10.35 | 12.91 | 9.360 | 52.64 | 29.31 | 31.77 | 30.16 | 48.96 | 30.75 | 41.18 | 26.71 |
| P-value | .0001 | .0001 | .0001 | .0001 | .0001 | .0001 | .0001 | .0001 | .0001 | .0001 | .0001 | .0001 | .0001 | .0001 | .0001 | .0001 |
| $\eta^2$ | .0165 | .0122 | .0149 | .0106 | .0096 | .0093 | .0059 | .0046 | .0116 | .0116 | .0138 | .0131 | .0095 | .0124 | .0150 | .0110 |
| **Error** | .4437 | .4768 | .4768 | .5626 | .3275 | .6759 | .6509 | .6782 | .3030 | .5954 | .5949 | .5719 | .2636 | .5508 | .4899 | .5633 |

*Sex* differences in WM variability have also been well-documented, with females generally exhibiting lower global FA values compared to men **[20]**. In this study (**Fig. 3**), global WM integrity also followed documented behaviour, demonstrating lower global FA values in females (0.373 ± 0.095) compared to males (0.374 ± 0.093). Interestingly, similar AD values (AD = 0.001) were found in females (± 3.08 × 10-4) and males (± 2.93 × 10-4), whereas in both MD and RD, females (8.31 × 10-4 ± 2.51 × 10-4; 6.70 × 10-4 ± 2.36 × 10-4, respectively) had higher values than males (8.15 × 10-4 ± 2.32 × 10-4; 6.53 × 10-4 ± 2.14 × 10-4, respectively). However, the effect of *Sex* is much less pronounced compared to *Age* effects alone. The 4-way ANOVA (**Table 3**) also demonstrates in this case, that there is a significant level of variability associated with *Sex* as a factor (all p < 0.001), albeit with a very small effect size (𝜂^2^ < 0.02). Principal component analysis (PCA) was also performed on FA values with *Sex* included as a factor across all *ROIs*. Interestingly, the results revealed that males (49.5%) and females (50.5%) contributed almost equally to the total variance, suggesting minimal *Sex*-related differences in global FA compared to other factors. Investigating *Sex* as a function of FA maps (**Fig. 4**) also demonstrates high levels of variability in the interhemispheric areas like the CC, a notable area affected by *Sex* differences.

**Figure 3.**
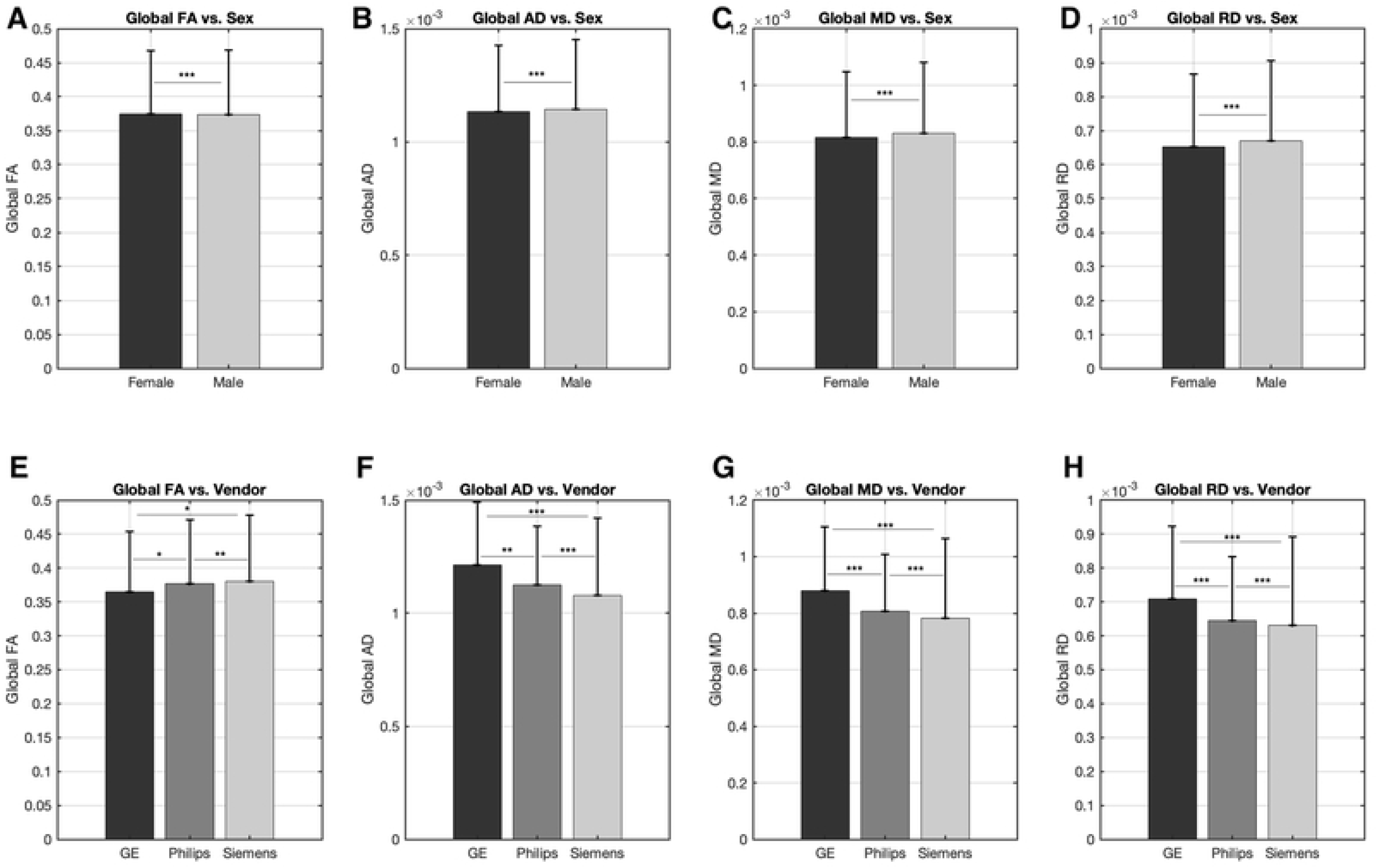
(A-D) Bar graphs illustrating values of global diffusion tensor imaging (DTI) measures pooling all ROIs and stratified by *Sex* (Male and Female). (A) Fractional Anisotropy (FA), (B) Axial Diffusivity (AD), (C) Mean Diffusivity (MD), and (D) Radial Diffusivity (RD) measures demonstrate how Female groups (black) have slightly lower global FA values and higher MD and RD values compared to males (light gray). Upper bound errors bars are also shown for each group and DTI metric and asterisks (*, **, ***) represent between group differences (𝑝 < 0.05, 𝑝 < 0.01, 𝑝 < 0.001, respectively). (E-H) Bar graphs depicting global diffusion measures pooling all regions of interest (*ROIs*) across three different *Vendors* in GE (black), Philips (dark gray), and Siemens (light gray). Similarly, (E) FA, (F) AD, (G) MD, and (H) RD values highlight variations introduced by *Vendor* with Siemens showing slightly higher values in FA and GE showing higher values in AD, MD, and RD. Upper bound errors bars are also shown for each *Vendor* and DTI metric and asterisks (*, **, ***) represent between group differences (𝑝 < 0.05, 𝑝 < 0.01, 𝑝 < 0.001, respectively).

**Figure 4.**
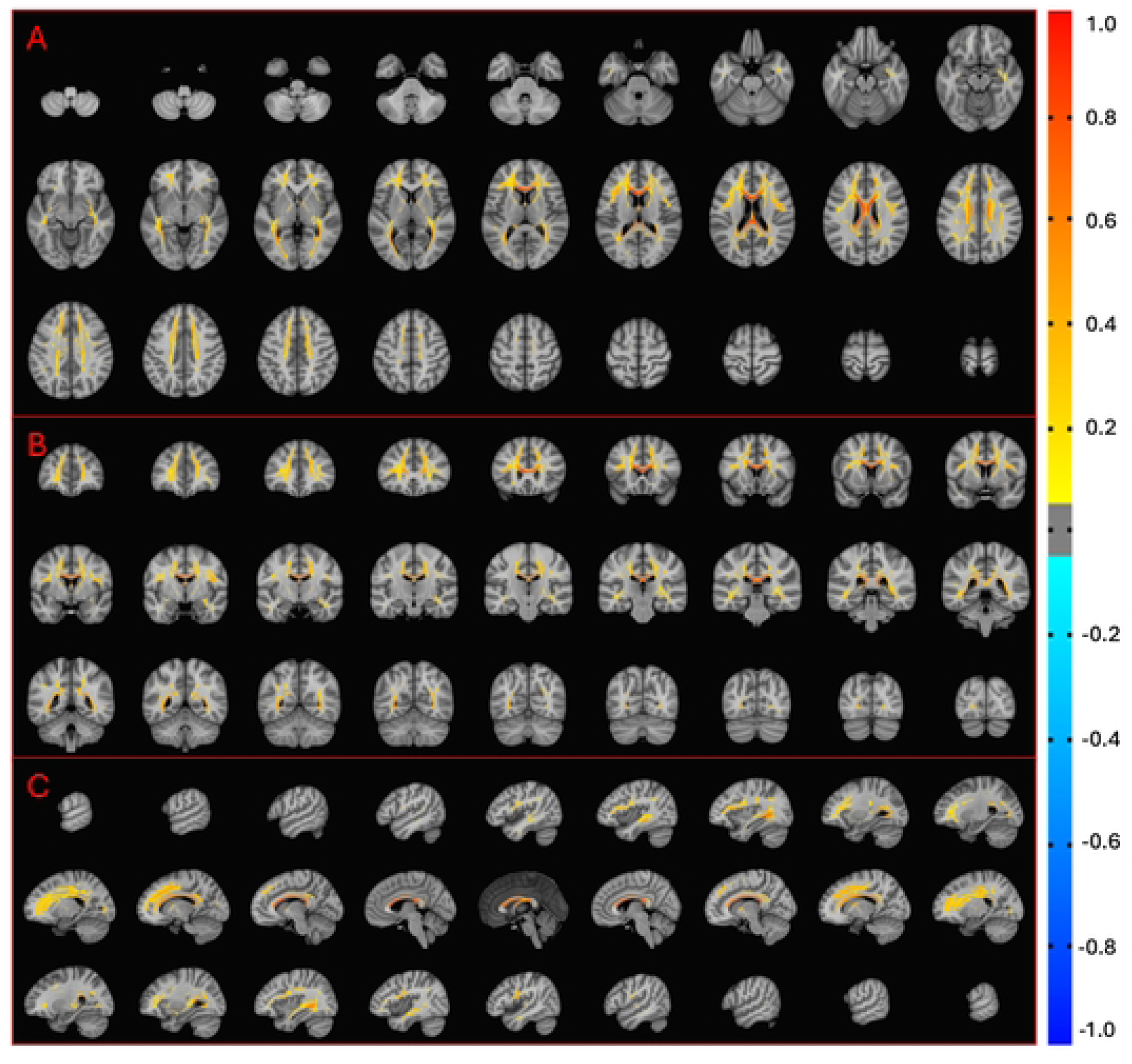
Variability in Fractional Anisotropy (FA) across the brain due to *Sex*, visualized using a 9 × 3 grid of axial (A), coronal (B), and sagittal (C) slices from a *3dANOVA* analysis in AFNI **[66]**. Analysis was performed on FA maps of all individuals (𝑛 = 2700) stratified in Male or Female groups. Regions with higher FA variability are shown in orange, particularly concentrated near interhemispheric areas such as the corpus callosum (CC), indicating pronounced *Sex*-related changes in these areas. Yellow regions represent significant but less extreme variability, reflecting more moderate *Sex*-related alterations. This visualization highlights the heterogeneous effects of *Sex* on white matter (WM) integrity across different brain regions and demonstrates that *Sex* has less overall variability effects compared to *Age*-related changes.

Previous literature has also shown that *Vendor* differences exist, particularly in MD and RD values **[26]**, with specific manufacturers like Siemens scanners exhibiting higher reliability than GE or Philips machines **[24]**. In this study (**Fig. 3**), global FA values for Siemens (0.380 ± 0.098) were also proven to be higher than Philips (0.377 ± 0.094) and GE (0.365 ± 0.088) machines. Interestingly, the FA error values were smaller in GE machines compared to both Siemens and GE, showing even more technical nuances related to *Vendor* behaviour. Comparatively, inverse relationships were seen for AD MD, RD with GE having higher values compared to Siemens and Philips. The 4-way ANOVA (**Table 3**) also demonstrated, in this case, that there is a significant level of variability associated with *Vendor* as a factor, with some medium effect sizes (𝜂^2^ > 0.06) in the JHU-Tracts *Atlas*. PCA was also performed on FA values across all *ROIs* with *Vendor* included as a factor. The results identified that Siemens (30.5%) had considerably less contributions to variance than Philips (34.7%) and GE (34.8%). *Vendor* as a function of FA maps (**Fig. 5**) also showed high levels of variability but in areas more associated with geometric distortions and signal dropout, predominantly near the ventricles.

**Figure 5.**
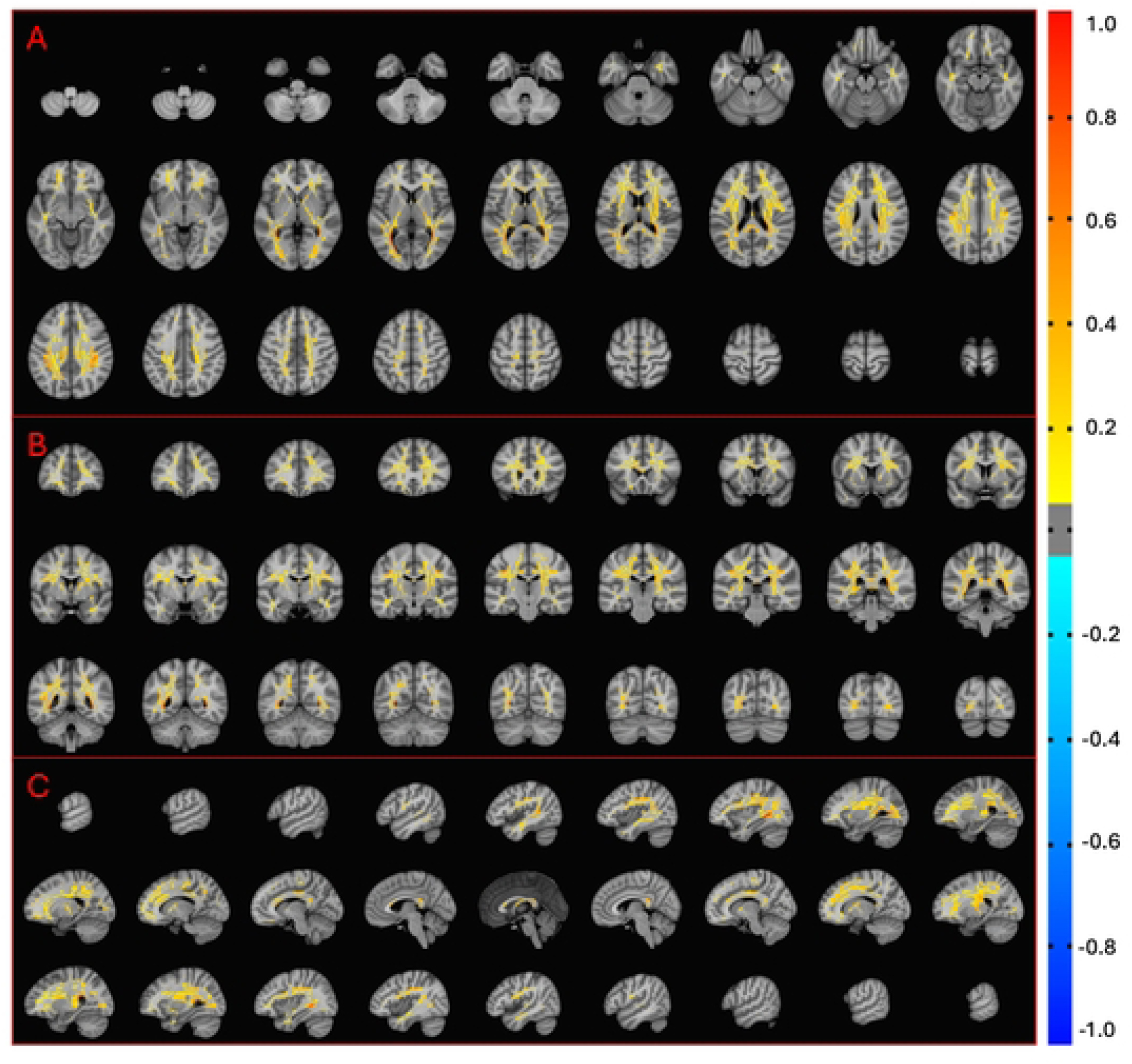
Variability in Fractional Anisotropy (FA) across the brain due to *Vendor*, visualized using a 9 × 3 grid of axial (A), coronal (B), and sagittal (C) slices from a *3dANOVA* analysis in AFNI **[66]**. Analysis was performed on FA maps of all individuals (𝑛 = 2700) stratified in their respective GE, Siemens, Philips *Vendor* groups. Regions with higher FA variability are shown in orange, particularly concentrated near the ventricles, indicating pronounced *Vendor*-related changes in these areas. Yellow regions represent significant but less extreme variability, reflecting more moderate *Vendor*-related alterations. This visualization highlights the heterogeneous effects of *Vendor* on white matter (WM) integrity measurements and demonstrates that *Vendor* has less overall variability effects compared to *Age*-related changes.

*ROIs* also have notable FA variability (1-4.2% change) and specific tracts are known to exhibit high variability in areas such as the UF (3.7-6% change) **[59]**. *ROI* variability has already been shown in this study where interhemispheric differences were found due to *Age* and *Sex* effects, while the edges of the ventricular areas are highly variable due to *Vendor*-related effects. *ROI* variability can also be associated with specific tract functions such as in sensory-motor tracts like the corticospinal tract (CT) due to late-stage neural development and in higher-order association areas (i.e. frontal tracts) that regulate cognition and emotion **[22]**. This study also focused on choosing multiple *Atlases* to investigate the differences between ROI selections. This is highlighted in the 4-way ANOVA (**Table 3**) that demonstrates each *Atlas* and all *ROIs* have significant variability with large effect sizes in the JHU-Tracts atlas. However, PCA on FA variability (**Fig. 6**) showed that JHU-Tracts *ROIs* have low variance contributions compared to JHU-Labels *ROIs*; whereas some XTRACT *ROIs* had low variance like the acoustic radiation (AR) and inferior longitudinal fasciculus (ILF), while the cingulum perigenual (CP) had high variance. It should also be noted that the JHU-Labels tapetum of the CC had the highest contribution of variance in this dataset.

**Figure 6.**
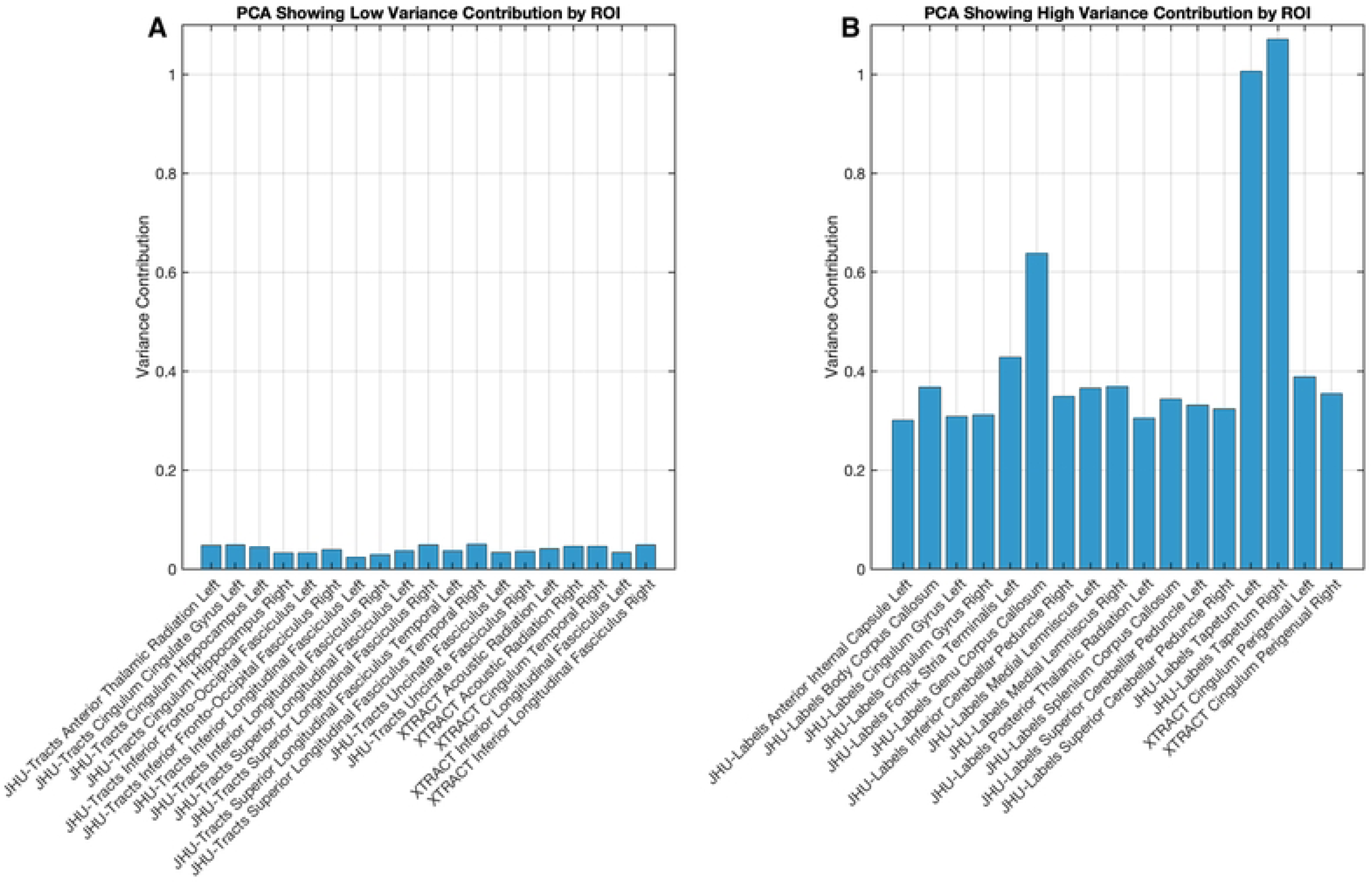
Principal component analysis (PCA) results illustrating the variance contribution of fractional anisotropy (FA) values across regions of interest (*ROIs*). (A) *ROIs* with low variance contributions (below the threshold of 0.05), indicate areas with minimal variability across the cohort. (B) *ROIs* with high variance contributions (above the threshold of 0.30), highlighting regions with substantial impact on data variability. Variance contributions were calculated for each *ROI* after grouping data by *Age*, *Sex*, *Vendor*, and standardizing the data to zero mean and unit variance. Less variability is shown in JHU-Tracts *ROIs* (A) compared to JHU-Labels (B).

### Interaction Effects of Age and Regions of Interest on Fractional Anisotropy

*Age*-related effects on global FA variability further extend into quantifiable *Age*-related interaction effects within certain *ROIs*. For example, the CC and anterior/posterior cingulum have been previously linked to *Age*-related changes, explained by a loss of executive function **[20,21]**.

Comparatively, the CT is known to be more resistant to *Age*-related changes when preservation of motor function remains intact **[65]**. In this study, a heatmap helps visualize the effects of *Age* on FA values across JHU-Labels (**Table 4**). Most *ROIs* show a consistently small decline in FA as people healthily *Age*, with some *ROIs* showing large decreases such as the tapetum/body/genu of the CC and the posterior thalamic radiation (PTR). Surprisingly, a slight increase in FA was observed in the cingulum gyrus and superior cerebellar peduncle. Another heatmap (**Table 5**), outlines the effects of *Age* on FA values across *ROIs* derived from the JHU-Tracts and XTRACT *Atlases*. JHU-Tracts demonstrated consistently small declines across all *ROIs* with the CT exhibiting the highest FA values among *ROIs*. The XTRACT *Atlas* also showed consistently small declines across *ROIs*, with larger declines in the arcuate fasciculus (AF), forceps (minor/major), optic radiation (OR), and the thalamic radiation. In the XTRACT *Atlas* however, the cingulum temporal/dorsal/perigenual regions remained relatively consistent across *Age*, contradicting some of the findings in the JHU-Labels *Atlas*. The 4-way ANOVA results (**Table 3**) illustrate that *Age-ROI* interactions are all significant, albeit with none to small effect sizes for the JHU-Tracts and XTRACT *Atlases* only. Another heatmap (**Table 6**) displays the coefficients of variation (CVs) associated with FA values and *Age* across WM *ROIs* in the JHU-Labels *Atlas*. The CVs were revealed to increase with *Age*, with *ROIs* exhibiting the highest variability in the 60-65-year-old group, particularly the gyrus/hippocampus of the cingulum, the terminal fornix stria (FST), the tapetum/genu of the CC, and the superior fronto-occipital fasciculus (SFOF). Notable *ROIs* that show little variation across *Age* include the cerebral peduncle, retrolenticular internal capsule, superior corona radiata (SCR), and the superior longitudinal fasciculus (SLF). Another heatmap (**Table 7**) shows the CVs for the JHU-Tracts and XTRACT *Atlases*. In this case, the JHU-Tracts *ROIs* show similar levels of variability across *Age* compared to the XTRACT *Atlas* where most *ROIs* show an increase in variability due to *Age*. However, some ROIs have a highly acute level of variability in the younger *Age* groups, particularly in the XTRACT *Atlas* in the anterior commissure (AC), anterior thalamic radiation (ATR), and CP. Interestingly, the CVs are much smaller in these *Atlases* compared to the JHU-Labels results. A circular bar plot (**Fig. 7**) also demonstrates the mean and standard deviation differences across younger and older *Age* groups (20-25 age group in blue, and 60-65 age group in orange). The outer circle illustrates mean FA values for all *Atlas ROIs* with notable trends highlighting a decrease in FA in older *Age* groups in the JHU-Labels *Atlas*, particularly in the genu/body of the CC, PTR, and fornix ROIs. JHU-Tracts and XTRACT *Atlases* showed overall reduction in FA due to *Age* across most *ROIs*. Interestingly, all cingulum-related areas across *Atlases* showed either no difference or a slight increase in mean FA values due to *Age*, which again demonstrates the different behaviors *ROIs* can exhibit across the aging spectrum. The inner circle shows standard deviation differences across younger and older *Age* groups with variability increasing with *Age*. Notable increases in standard deviation due to *Age* are manifested in the XTRACT cingulum perigenual and anterior commissure, along with the JHU-Labels medial lemniscus, inferior fronto-occipital fasciculus (IFOF), peduncle related areas, cingulum related areas, and the CT.

**Table 4.**
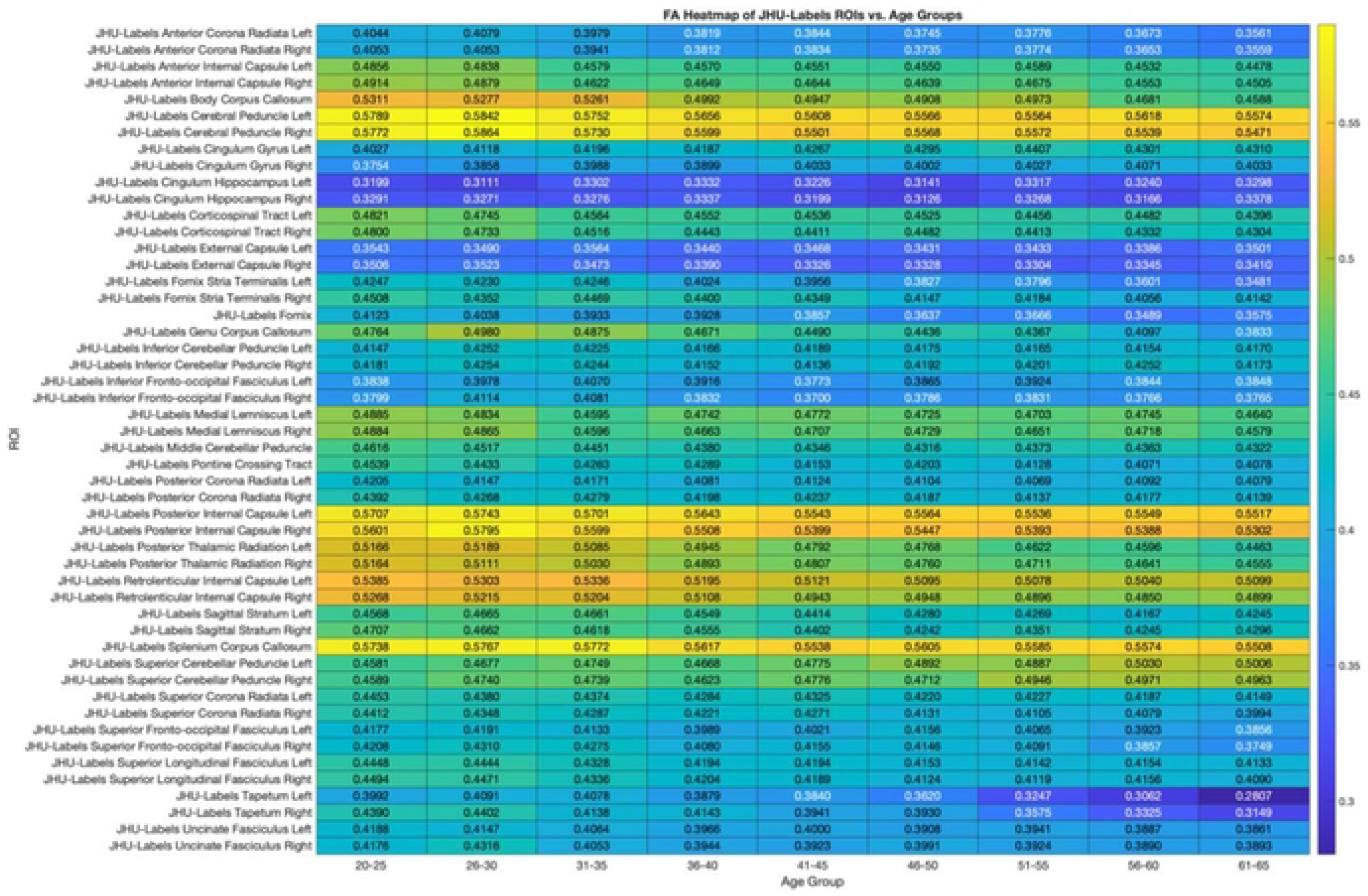
Heatmap visualizing mean fractional anisotropy (FA) values across 50 regions of interest *(ROIs)* from the JHU-Labels *A Has,* stratified by nine 5-year *Age* groups ranging from 20-65 years. Each cell represents the mean FA value for a specific *ROI* within an *Age* group, with warmer colors (yellow) indicating higher FA values and cooler colors (blue) indicating lower FA values. FA decreases with *Age* across most *ROIs* reflecting/Ige-related white matter (WM) changes. Certain tracts, such as the corpus callosum (CC) exhibit notable variability in FA across *Age,* potentially highlighting areas sensitive to aging effects.

**Table 5.**
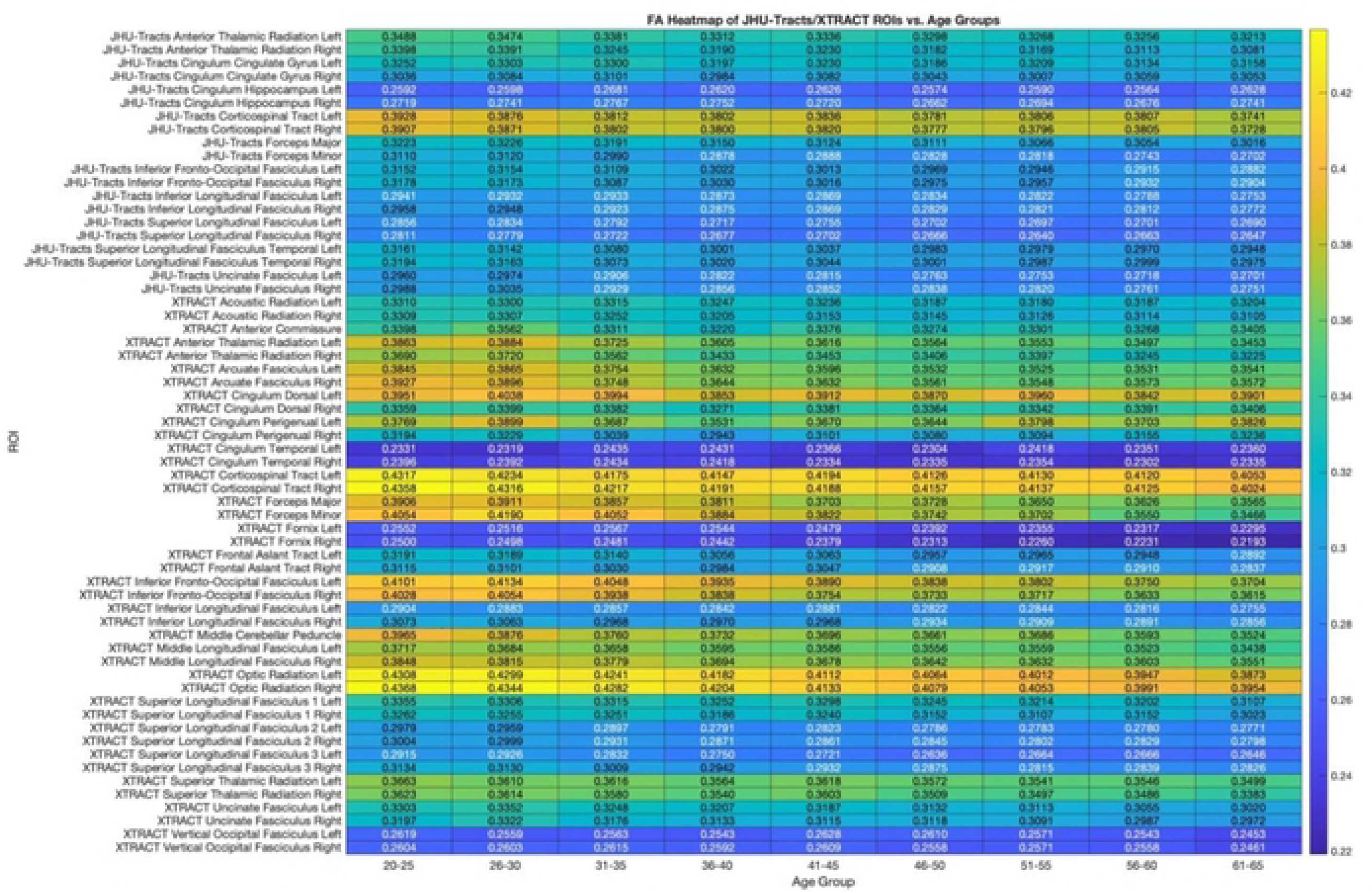
Heatmap showing mean fractional anisotropy (FA) values across 62 regions of interest *(ROIs)* from the JHU-Tracts (20) and XTRACT (42) *Aliases* across nine 5-year *Age* groups ranging from 20-65 years. Color intensities represent FA values with warmer colors (yellow) indicating higher FA values, while cooler colors (blue) represent lower values. Certain *RO/s* demonstrate distinct patters of FA changes associated with *Age,* while others show relatively stable FA values across *Age* groups. Pronounced declines are more often observed in association tracts compared to motor tracts. Differences across *Age* groups provide insights into *Age*-dependent microstructural changes.

**Table 6.**
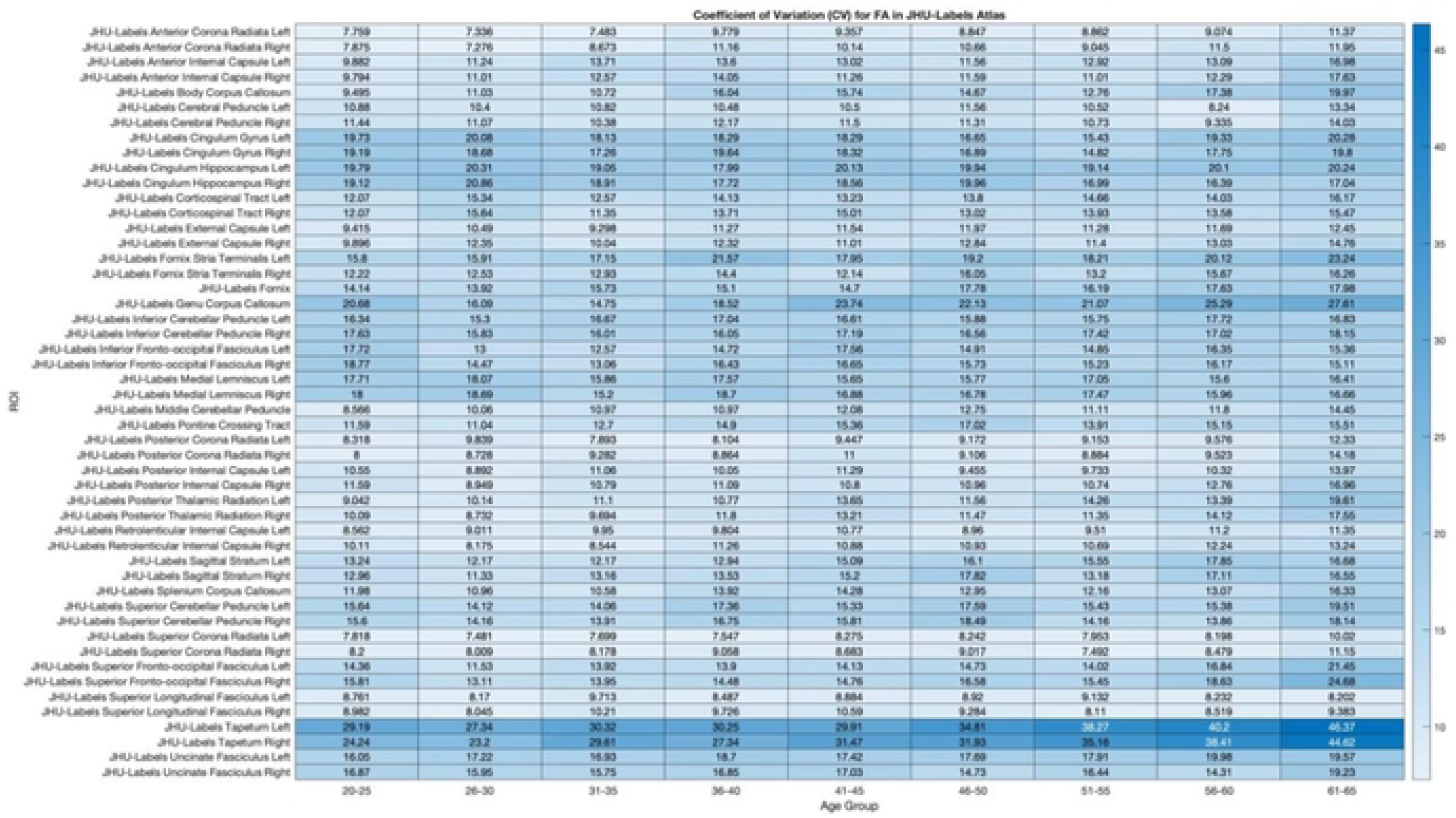
The heatmap displays the Coefficient of Variation (CV) of fractional anisotropy (FA) across 50 regions of interest *{ROIs)* from the JHU-Labels *Allas,* stratified by nine 5-year *Age* groups ranging from 20-65 years. CV is a measure of variability, normalized by the mean, where lower CV values (white) indicate more consistent FA measurements across *Age,* while higher CV values (dark blue) indicate greater variability. CV was computed as the ratio of the standard deviation to the mean of FA values within each group. Notably, the anterior/superior corona radiata (ACR/SCR) exhibit relatively low CV values (< 10%) across all *Age,* indicating stability. In contrast, regions such as the fomix and tapetum of the corpus callosum (CC) demonstrate higher CV values (> 20%), suggesting greater variability in FA. This variability underscores the regional differences in white matter (WM) integrity that are susceptible to aging.

**Table 7.**
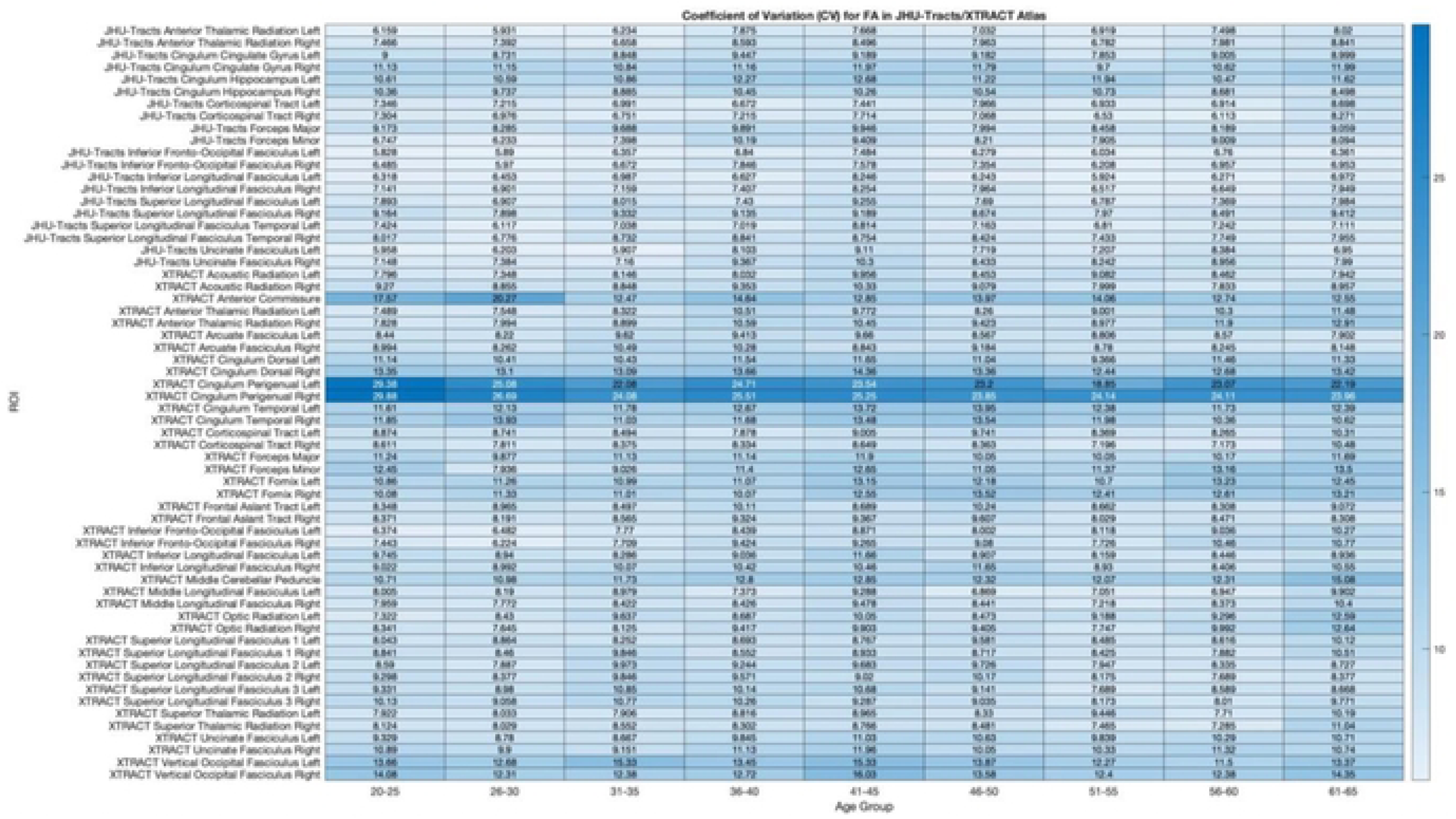
This heatmap illustrates the Coefficient of Variation (CV) for fractional anisotropy (FA) across 62 regions of interest (*ROIs*) from the JHU-Tracts (20) and XTRACT (42) *Aliases* across nine f year *Age* groups ranging from 20-65. Regions with lower CV values (white) exhibit more consistent FA measurements and indicate lower variability (greater consistency), while higher CV values (dark blue) indicate greater variability (less consistency). Within the JHU-Tracts *A lias,* the anterior thalamic radiation (ATR) and uncinate fasciculus (UF) demonstrate relatively low CV values, highlighting consistent white matter (WM) integrity through healthy aging. Conversely, in the XTRACT *Allas,* the cingulum gyrus/dorsal/temporal show increased CV values, suggesting greater heterogeneity. The variability in CV in *ROIs* may reflect differential susceptibility in WM integrity due to *Age*.

**Figure 7.**
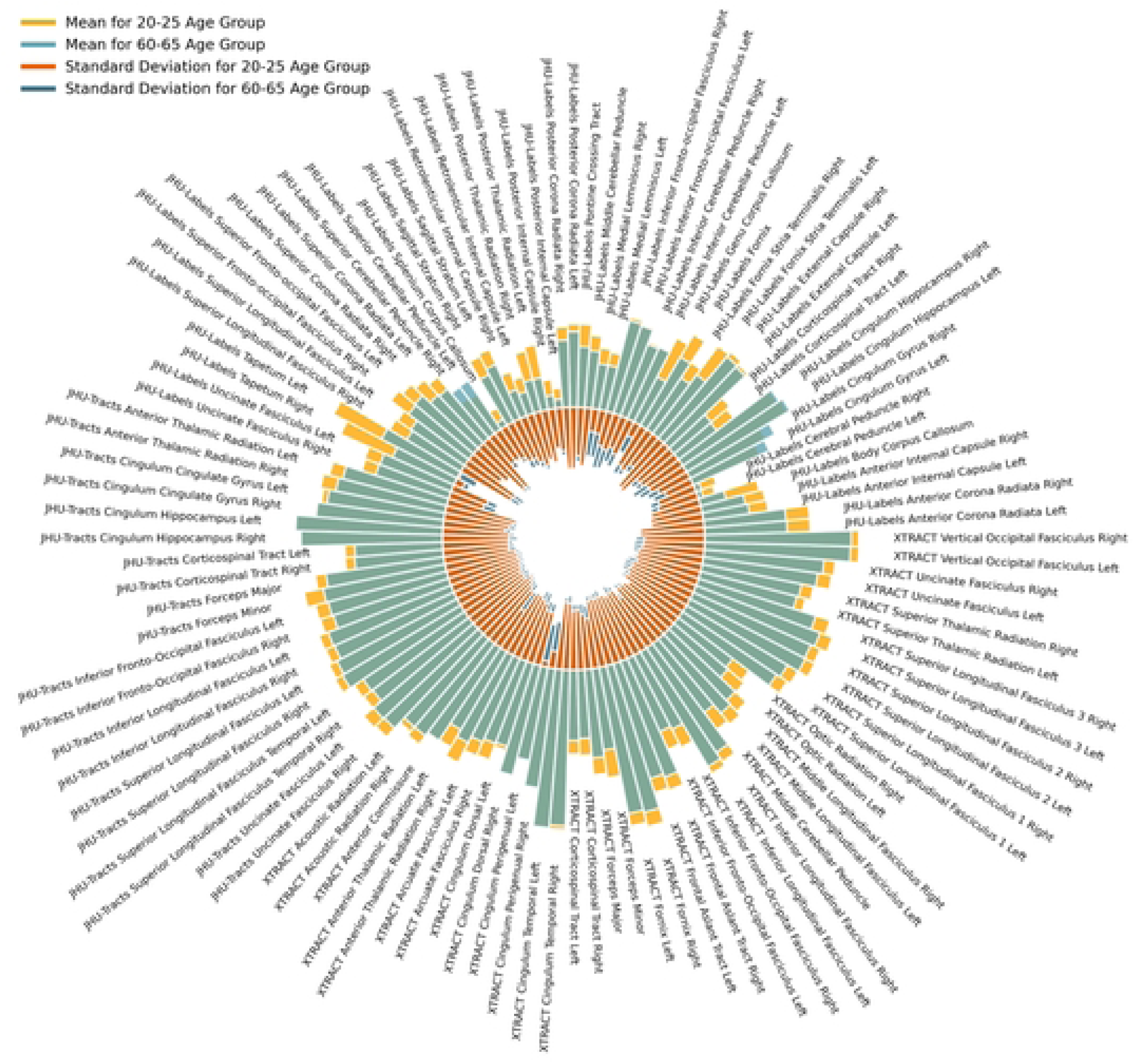
Comparison of mean Fractional Anisotropy (FA) values and standard deviations for two *Age* groups: 20-25 years and 60-65 years. The outer radial plot illustrates the mean FA values for each region of interest (*ROI*) for all *Atlases* and for 20-25 year old (light blue) and 60-65 year old (light orange) *Age* groups. The inner radial plot shows the mean standard deviation values for FA associated with each *ROI* and younger (dark blue) and older (dark orange) *Age* groups. Notable trends in the data include higher mean FA values and lower standard deviation values in the 20-25 year old *Age* group across most *ROIs*, indicating better and more consistent white matter (WM) integrity in younger individuals. These findings highlight the eclectic *Age*-related changes in WM organization across various brain regions.

### Interaction Effects of Sex and Regions of Interest on Fractional Anisotropy

*Sex*-related differences in *ROIs* on FA values highlight distinct patterns of WM structure and maturation. Historically, it has been shown that women tend to demonstrate lower global FA and higher MD values compared to men, particularly in the anterior cingulum and SLF and, with aging, this is often linked to metabolic changes due to post-menopause **[20]**. Comparatively, men often exhibit increases in FA and decreases in RD values compared to women, particularly in the CC and in association tracts, theorized to be associated with motor abilities and estrogen’s protective effects in women **[65]**. In this study, a heatmap helps visualize the effects of *Sex* on FA values across JHU-Labels, JHU-Tracts, and XTRACT *Atlases* (**Table 8**). Most *ROIs* show congruency between *Sex*, however, in the JHU-Labels *Atlas*, notable reductions in FA in females were found in the PTR and tapetum of the CC. Interestingly, females had higher FA in the external capsule compared to males. In other *Atlases*, no notable trends were found between *Sex*. An interaction line plot (**Fig. 8**) helps visualize the differences in mean FA values across all *ROIs* associated with *Sex*-differences. This plot also supports how different *Atlases* behave with larger differences highlighted in the JHU-Labels *Atlas*, particularly in the PTR and tapetum of the CC compared to small differences in the JHU-Tracts/XTRACT ANOVA results (**Table 3**) show that *Sex*-*ROI* interactions have significant variability across *ROIs*, albeit with small to no effect sizes. Another heatmap (**Table 9**) illustrates the CVs associated with FA values and *Sex* across WM *ROIs* in the JHU-Labels, JHU-Tracts, and XTRACT *Atlases*. Most cases demonstrate an overall increase in variability associated with females compared to males. In the JHU-Labels *Atlas*, it is shown that females have particularly high FA variability compared to males in the CT, FST, tapetum/genu of the CC, PTR, and the SFOF. Conversely, the JHU-Tracts *Atlas* showed relative consistency across male and female cohorts, whereas the XTRACT *Atlas* highlighted the CP, forceps minor/major, middle cerebellar peduncle, and OR as *ROIs* with higher FA variability in females. Interestingly, the XTRACT *Atlas* AC had higher FA variability in males.

**Figure 8.**
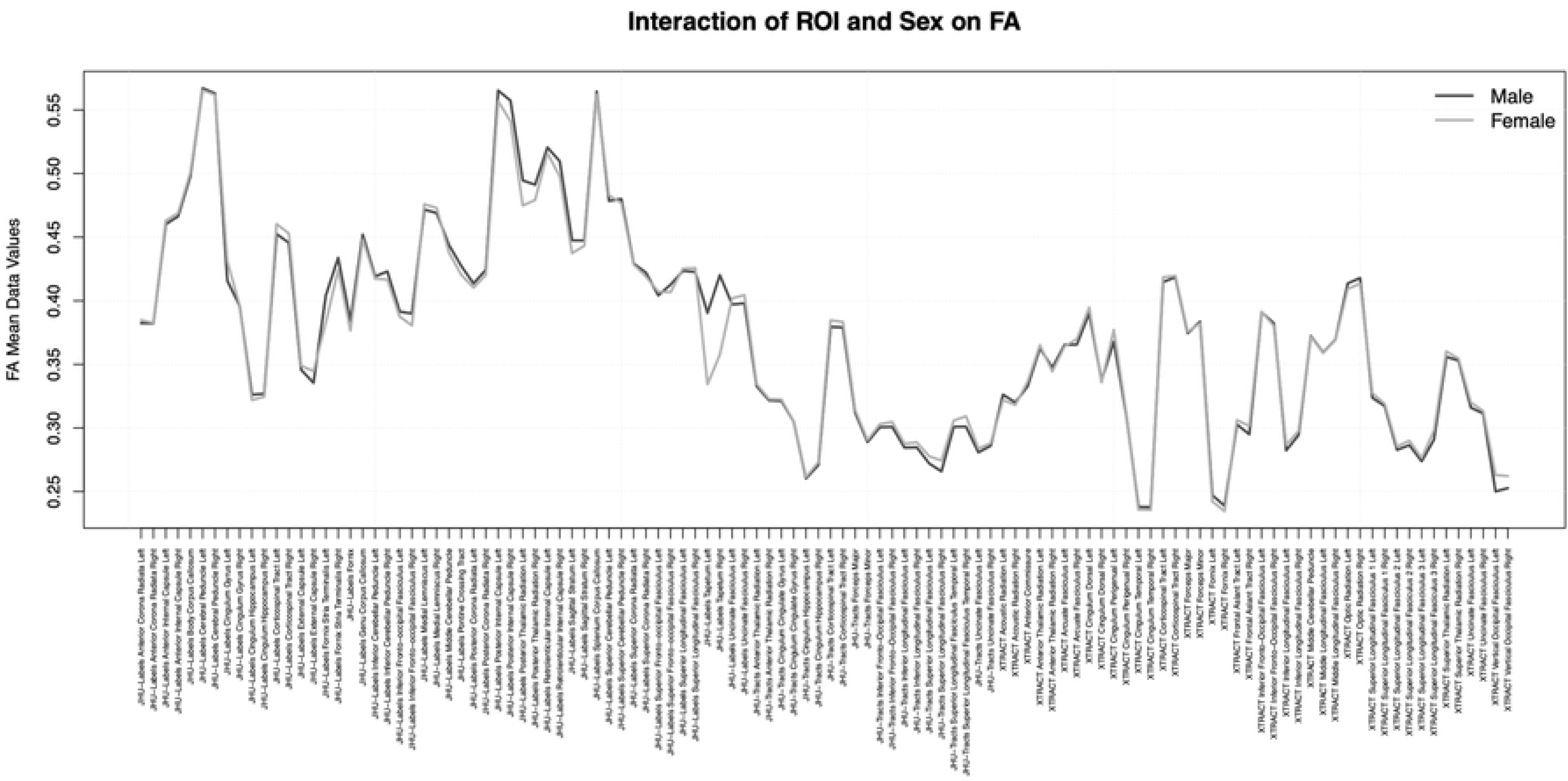
Interaction of region of interests (*ROIs*) and *Sex* on fractional anisotropy (FA) mean values in all three *Atlases* (JHU-Labels, JHU-Tracts, XTRACT). This line graph compares the mean FA values for male (dark gray) and female (light gray) subjects, providing insights into the variability of white matter (WM) integrity across different brain structures. Male and female FA values demonstrate similar trends yet in some select *ROIs*, males tend to exhibit higher FA values in the JHU-Labels posterior thalamic radiation (PTR) and the tapetum of the corpus callosum (CC), suggesting potential differences in WM organization. Notable fluctuations in FA across *ROIs* also indicate substantial variability due to *Sex*, underlining the complex interplay between biological *Sex* and brain structure.

**Table 8.**
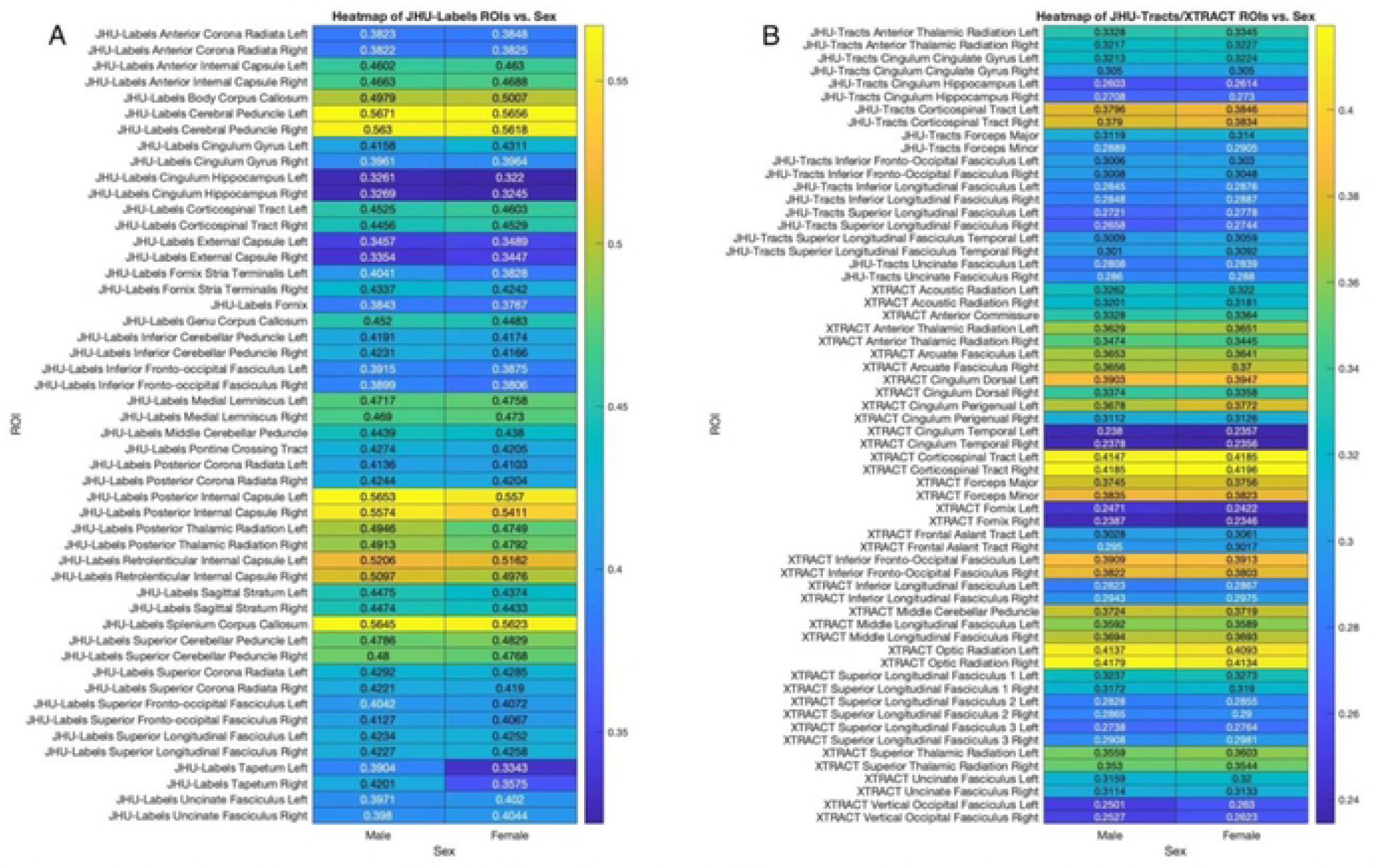
A pair of heatmaps illustrating Sex-based differences in mean Fractional Anisotropy (FA) values across white matter (WM) regions of interest *(ROIs)* in the JHU-Labels *Allas* (A) and the JHU-Tracts/XTRACT *Aliases* (B). The color scale represents the intensity of mean FA values where wanner colors (yellow) indicate higher values compared to cooler colours (blue) representing lower values, showing the differences between *Sex.* Most *ROIs* show consistency in FA values across *Sex,* however, key *ROIs* in the JHU-Labels *Allas* such as the fornix stria terminalis (FST), posterior thalamic radiation (PTR), and tapetum of the corpus callosum (CC) show that females have lower mean FA values compared to males. In contrast, the JHU-Tracts and XTRACT *Allas* show more consistency across *Sex,* with minute differences between the two groups. These results emphasize the distinct patters of WM microstructure variability linked to *Sex*.

**Table 9.**
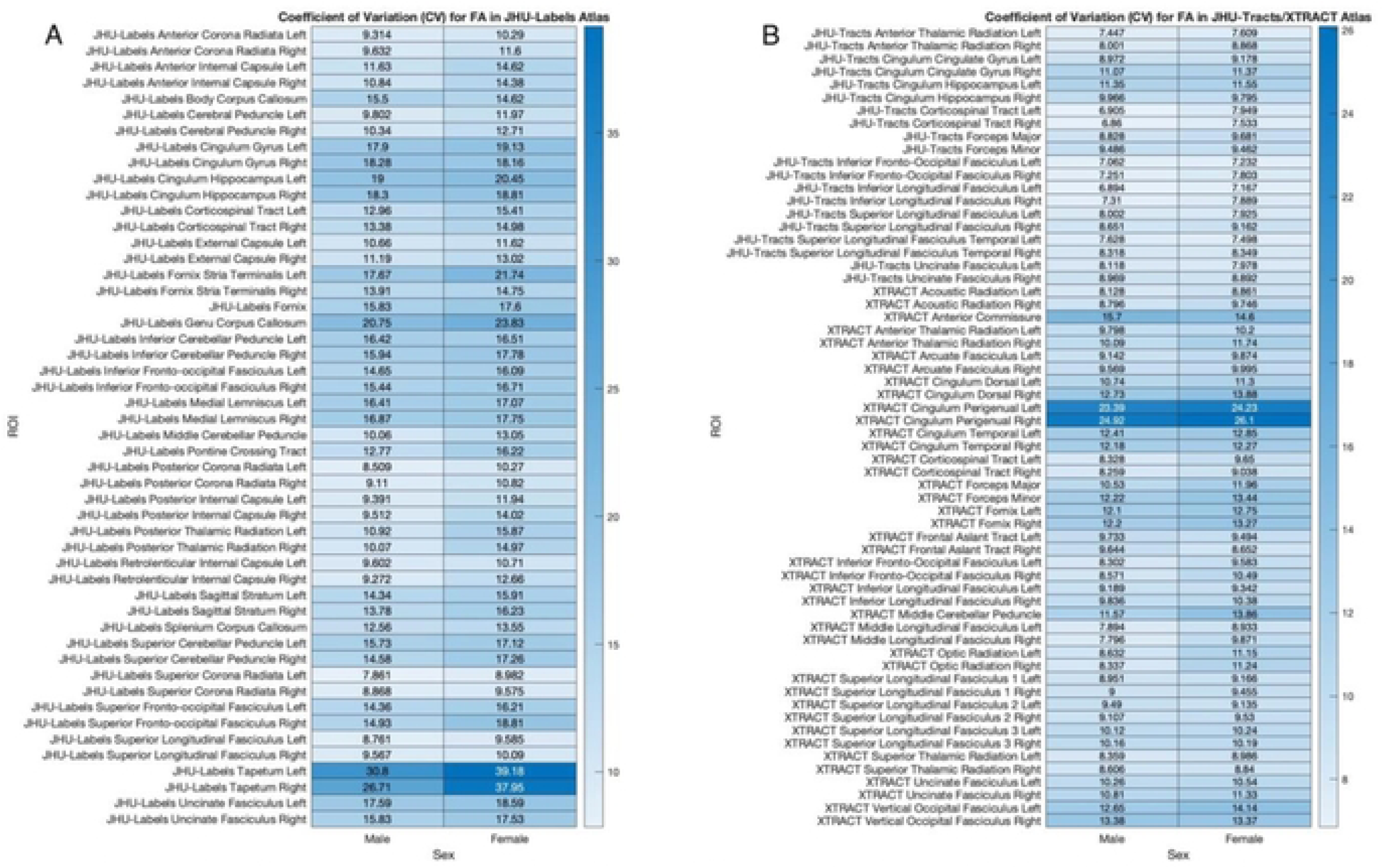
A pair of heatmaps showcasing the coefficient of variation (CV) table for fractional anisotropy (FA) associated in various regions of interest *(ROIs)* across male and female subjects, utilizing the JHU-Labels *Allas* (A) and the JHU-Tracts/XTRACT *Aliases* (B). Color intensities depict lower CV values (white) having greater consistency compared to *ROIs* with higher CV values (dark blue) having greater variability. The data reveals notable differences in CV, indicating variability in FA measurements across different brain structures due to *Sex.* Specific trends include lower CV values in the superior longitudinal fasciculus (SLF) and superior corona radiata (SCR) in the JHU-Labels *Allas* along with the corticospinal tract (CT), inferior fronto-occipital fasciculus (IFOF), and inferior longitudinal fasciculus (ILF) in the JHU-Tracts *A lias,* demonstrating stability in these *ROIs* across/Ige. Conversely, the fornix stria terminalis (FST), and the genu/tapetum of the corpus callosum (GCC/TCC), of the JHU-Tracts *Atlas* along with the cingulum perigenual (CP) of the XTRACT *Allas* show higher CV values and in turn greater variability differences between *Sex.* These findings highlight the importance of anatomical region and *Sex* interactions in the assessment of FA, with implications for improving the understanding of white matter (WM) integrity across various populations.

### Interaction Effects of Vendor and Regions of Interest on Fractional Anisotropy

*Vendor*-related differences in FA from *ROIs* are also an important aspect to investigate with regards to WM variability. Historically, variations in FA measurements have been linked to differences in scanner hardware and software configurations with Siemens exhibiting higher overall FA values compared to GE and Philips **[14,24]**. These scanner *Vendor* differences often manifest in specific *ROIs*, surrounding tissue susceptibility areas (i.e. proximal to the lateral ventricles) (**Fig. 3**). In this study, a heatmap (**Table 10**) visualizes the effects of *Vendor* variability on FA values across the JHU-Labels, JHU-Tracts, and XTRACT *Atlases*. Notable discrepancies were observed in the JHU-Labels *Atlas* for the cingulum, CT, cerebellar peduncle, and medical lemniscus ROIs, all of which showed the Siemens MRI scanners having the highest mean FA values. Conversely, the genu/body/splenium of the CC all had notably high FA values for Philips systems. Investigating the JHU-Tracts *Atlas*, the CT had much lower FA values in GE machines, while the forceps minor/major and the SLF had higher FA values in Philips machines. In the XTRACT *Atlas*, a variety of trends were found such that for Siemens, the AC, cingulum perigenual/temporal had higher FA values while the vertical occipital fasciculus had lower FA values compared to other systems. Moreover, in data from Philips scanners, the fornix had lower FA values while the ATR, arcuate fasciculus (AF), forceps minor/major, middle cerebellar peduncle, middle longitudinal fasciculus, OR, and UF all had higher FA values. The GE scanners tended to have lower FA values in both the CT and superior thalamic radiation (STR). A corresponding interaction plot (**Fig. 9**) provides a visual representation of the heatmaps (**Table 10**), where large differences between all three *Vendor* FA values are shown in all *ROIs*. *Vendor* emerged as the most consistent source of variability across *Atlas*, with similar FA differences observed in the CT, SLF, and UF across both the JHU-Tracts and XTRACT *Atlases*. Consistent with the heatmap findings, the cingulum dorsal and IFOF remain notably consistent across *Vendor* platforms. The results of the 4-way ANOVA (**Table 3**) for *Vendor*-*ROI* interactions exhibit significant variability with some small effect sizes. Another pair of heatmaps (**Table 11**) illustrates the CV associated with FA values across *Vendor*. For the JHU-Labels *Atlas*, higher CV values were routinely found in Philips machines, particularly in the cingulum gyrus/hippocampus, FST, tapetum/genu of the CC, medial lemniscus, and cerebellar peduncle *ROIs*. Comparatively, lower CV values were often found in GE and Siemens machines in *ROIs* such as the cerebral peduncle, SCR, and SLF. The JHU-Tracts *Atlas* showed similar patterns in that higher CVs were manifested in Philips machines in the cingulum gyrus/hippocampus and forceps minor/major *ROIs*, lower CVs were shown in GE machines in the CT, ILF, and SLF, while Siemens machines had low CVs in the UF. Similar patterns were repeated in the XTRACT *Atlas* with lower CVs found in the CT and SLF for GE and lower CVs in the UF for Siemens platforms.

**Figure 9.**
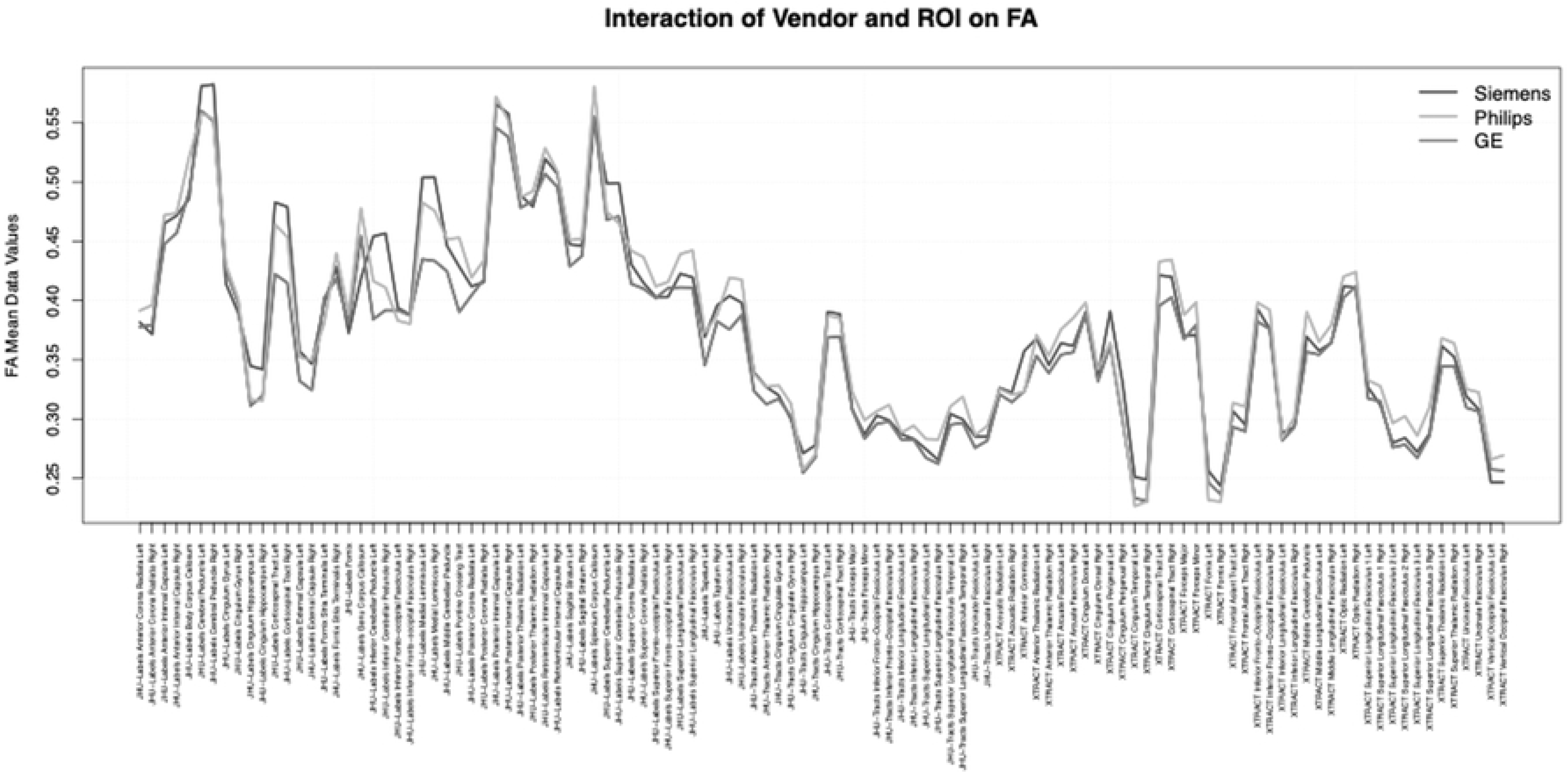
Plot line interaction display of *Vendor* (Siemens, GE, Philips) on fractional anisotropy (FA) mean values across all three *Atlases* (JHU-Labels, JHU-Tracts, XTRACT) to assess *Vendor* variability. Each line represents the mean FA values for a specific *Vendor* across *ROIs* where Siemens (black), GE (dark gray), and Philips (light gray) show fluctuating FA behaviour. In the JHU-Labels *Atlas*, Siemens has predominantly higher mean FA values, whereas in the JHU-Tracts/XTRACT *Atlases*, Philips tended to have higher FA values. These discrepancies were more pronounced in *ROIs* with higher inherent variability, such as the corticospinal tract (CT), the tapetum of the corpus callosum (CC), and uncinate fasciculus (UF). The variability observed in *ROIs* due to *Vendor*-specific differences highlights the need for careful consideration when using multi-site data in diffusion imaging studies.

**Table 10.**
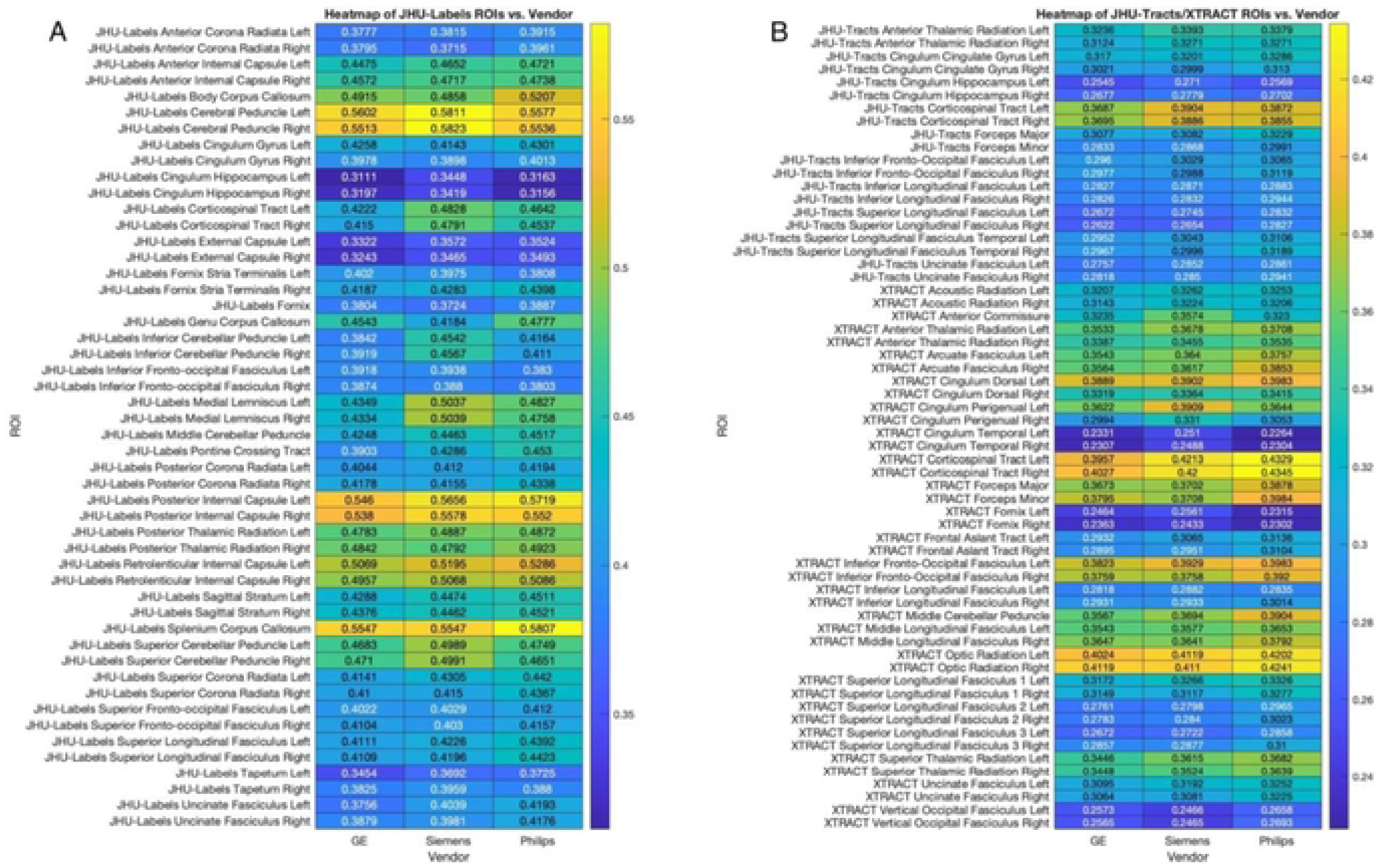
Pair of heatmaps showing fractional anisotropy (FA) values across MRI scanner *Vendors* (GE, Siemens, Philips) and regions of interest *(ROIs)* pertaining to the JHU-Labels *A f/as* (A) and the JHU-Tracts/XTRACT *Aliases* (B). Color intensities represent lower FA values (blue) and higher FA values (yellow) that highlight inter-FeWo/’ white matter (WM) sensitivities. Key differences across *Vendor* are evident in several *ROIs,* including the JHU-Labels *A das* in the cingulum, corticospinal tract (CT), cerebellar peduncle, medical lemniscus and all corpus callosum (CC) regions. In the JHU-Tracts *Atlas,* the CT, fornix minor/major and superior longitudinal fasciculus (SLF) were notably variable, whereas the XTRACT *Atlas* differences manifested in almost all ROIs. These findings underscore the influence of Fe/zt/w-related variability in FA measurements.

**Table 11.**
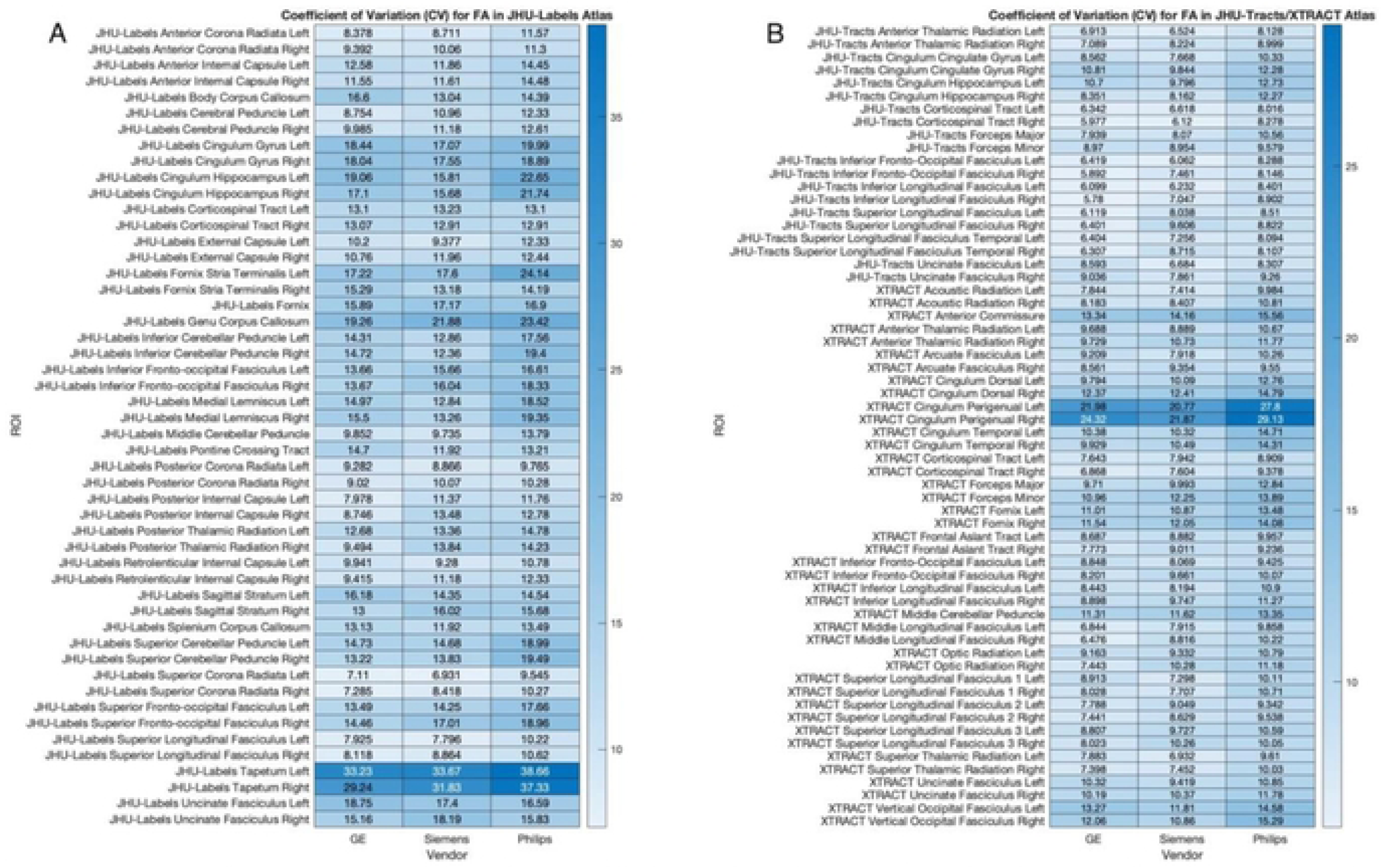
Two heatmaps showing the coefficient of variation (CV) for fractional anisotropy (FA) values in white matter (WM) across *Vendor* (GE, Siemens, Philips) and in various regions of interest *(ROIs)* selected from the JHU-Labels (A) and JHU-Tracts/XTRACT (B) *Atlases.* Each row represents a specific *ROI* with CV values indicated numerically and visually by color intensity with darker shades (dark blue) representing higher CV values. CV values were calculated as the ratio of standard deviation to the mean FA for each *Vendor ROI* combination. Notable differences in CV were observed across *Vendor* with the JHU-Labels tapetum of the corpus callosum (CC) and XTRACT cingulum perigenual (CP) demonstrating dark blue areas with particularly high *Vendor* variability. Regions such as the JHU-Labels superior corona radiata (SCR) and the XTRACT Superior Thalamic Radiation (STR) exhibited consistently low CVs, indicating more uniform FA measurements across *Vendor.* Moreover, the variability between *Vendor* in specific *ROIs* such as the XTRACT CP showed 10% differences in *ROI* measurements across *Vendor.* This analysis highlights the impacts of reproducibility of diffusion metrics across platforms.

**Table 12.** This covariance matrix (CM) displays the covariance values between different factors including *Age, Sex,* region of interest *(ROI),* and *Vendor.* This is a Hermitian matrix (symmetric) as the diagonal elements represent the variances for each factor, while off-diagonal elements indicate the covariance between factors. Notably, in bold, the variance for *ROI* is substantially larger compared to other factors, reflecting its high variability and suggesting strong internal consistency in measurements related to brain anatomy across participants. Covariance between factors is also small in comparison to the variance in *ROI*.

|  | *Age Group* | *Sex* | *ROI* | *Vendor* |
| --- | --- | --- | --- | --- |
| Age Group | 6.667 | -.0006 | .0002 | .0011 |
| Sex | -.0006 | .2500 | .0001 | .0002 |
| ROI | .0002 | .0001 | **1045** | .0001 |
| Vendor | .0011 | .0002 | .0001 | .6665 |

Alternatively, the CP, forceps minor/major, fornix, middle cerebellar peduncle, and vertical occipital fasciculus all had notably high CVs for Philips machines. These findings underscore the importance of accounting for *Vendor*-related differences in FA metrics, particularly in multicenter studies that integrate data from multiple scanning platforms.

### Interaction Effects of Age, Sex, Vendor, and Regions of Interest on Fractional Anisotropy

Interaction effects among all factors are also important to identify. It’s not always main effects in a statistical mode that fully determine the sources of variance in data. For instance, *Age*, *Sex*, *Vendor*, *Age*-*ROI*, *Sex*-*ROI*, and *Vendor*-*ROI* interactions have been discussed, but it is important to note in the 4-way ANOVA tables (**Table 3**), that *Age, Sex*, *Age*-*Vendor*, and *Sex*-*Vendor* contributions also exist. For example, *Age*-*Sex* interactions are significant, albeit with small to no effect sizes. **Fig. 10** shows that aging females exhibit an overwhelming increase in FA, while FA values in younger *Age* groups are more balanced between male and female. This also highlights the overall decrease in FA with *Age*. However, *Vendor* differences were also noted, with Siemens exhibiting higher FA values and GE exhibiting the lowest FA values among *Vendors*. *Sex*-*Vendor* interactions were similar to *Age*-*Sex* effects where they were calculated to have significance but with small to no effect sizes. These differences may be attributed to a correlation between head size and MRI-related artifacts, as males generally have larger heads than females, potentially affecting DTI fidelity through variations in signal-to-noise ratio, geometric distortions, and susceptibility artifacts **[15]**. A covariance matrix (CM) was also calculated (**Table 12**) to examine the interaction effects across factors. In this Hermitian style matrix (symmetric about the diagonal), it is clearly demonstrated that *ROIs* (1045) are the single largest contributor to variance, with *Age* also contributing significant variance (6.67). Interestingly, *Sex* (0.25) and *Vendor* (0.65) had much lower contributions within the covariance matrix. The CM also highlighted that interaction effects (off-diagonal symmetric cross-terms) were relatively negligible in comparison to *ROI* effects. The circular bar plot comparison, showcasing two cohorts of males and females in younger and older *Age* groups, helps articulate these nuances (**Fig. 11**). This also helps summarize important overall findings in that males generally have higher FA values than females, older females exhibit greater FA variability, and some *ROIs* have noteworthy behavioural characteristics. For instance, one of the most intriguing *ROIs*, the tapetum of the CC in the JHU-Labels *Atlas*, proves to be highly variational in both male and female cohorts, particularly in older *Age* groups. It should be expressed that each *ROI* must be scrutinized in a case-by-case basis as to understand the entire effects of *Age*, *Sex*, and *Vendor* differences associated with DTI measurements.

**Figure 10.**
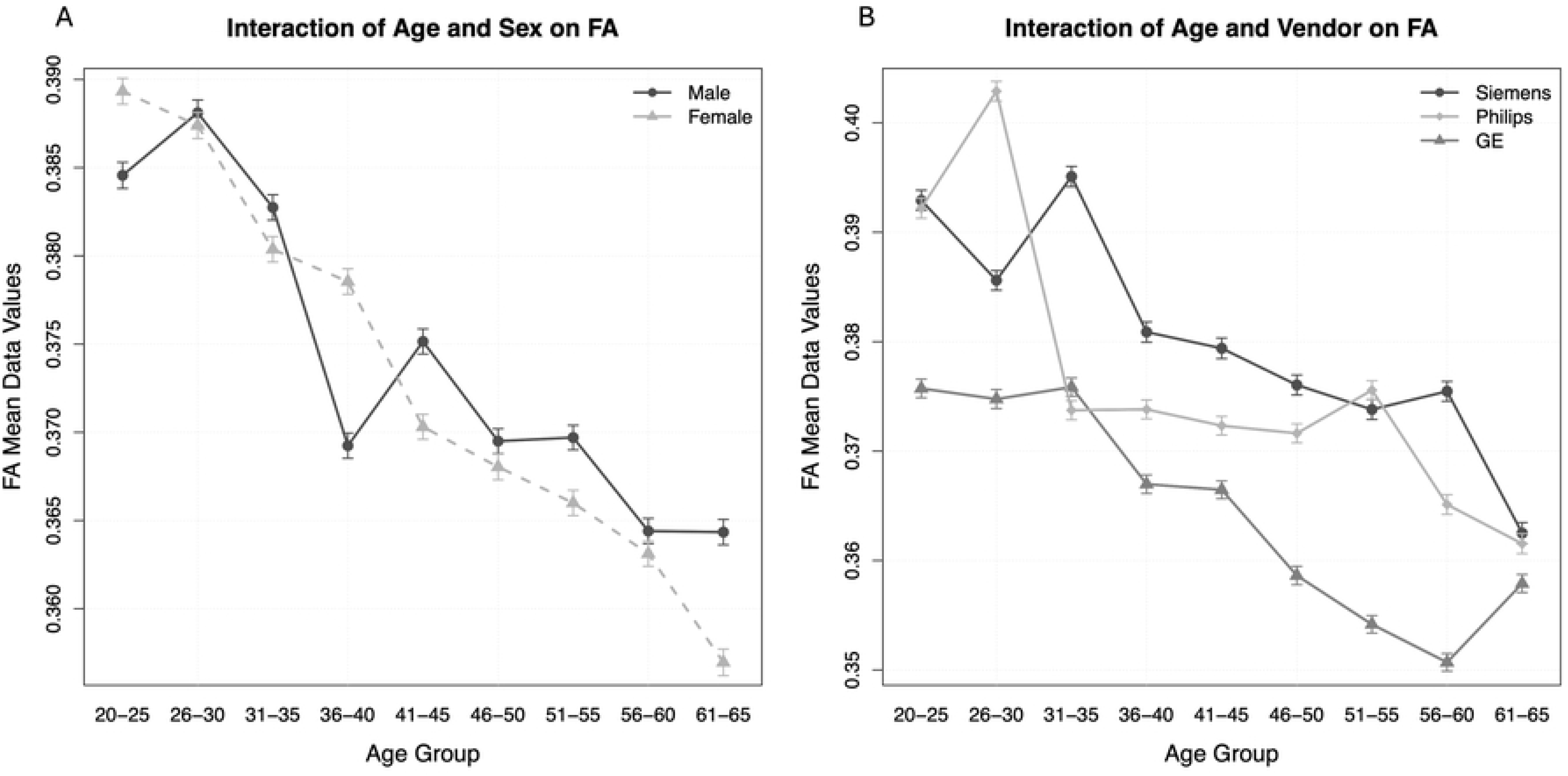
Pair of interaction plot lines that illustrate the interaction effects of *Age*-*Sex* and *Age*-*Vendor* on fractional anisotropy (FA) mean data values. For (A) *Age*-*Sex* interactions, males are represented by a solid black line and circular points while females are depicted by a dashed light gray line and triangular points, with error bars indicating standard error of the mean at each point. Both male and female subjects exhibited a decline in FA values across *Age* groups, while males consistently showed higher FA values than females, particularly in older *Age* groups (56-60 and 61-65 years). This indicates a potential *Sex*-specific difference in white matter (WM) integrity due to *Age*. For (B) *Age*-*Vendor* interactions, Siemens data is represented by a solid black line and circular points, Philips data by a light gray line and diamond points, and GE data by a dark gray line and triangular points, with error bars also indicating standard error of the mean at each point. Siemens data maintains higher FA values across most *Age* groups compared to Philips and GE platforms. Notably, the 31-35 *Age* group showed large disparities between Siemens and GE/Philips values, while the 51-55 year *Age* group shows a reduction in GE FA values compared to Siemens/Philips platforms. These findings suggest that *Vendor* differences might influence FA outcomes in different *Age* groups. Both plots illustrate trends in FA mean values across *Age*, revealing significant declines with aging and highlighting the influence of *Sex* and *Vendor* on FA measurements.

**Figure 11.**
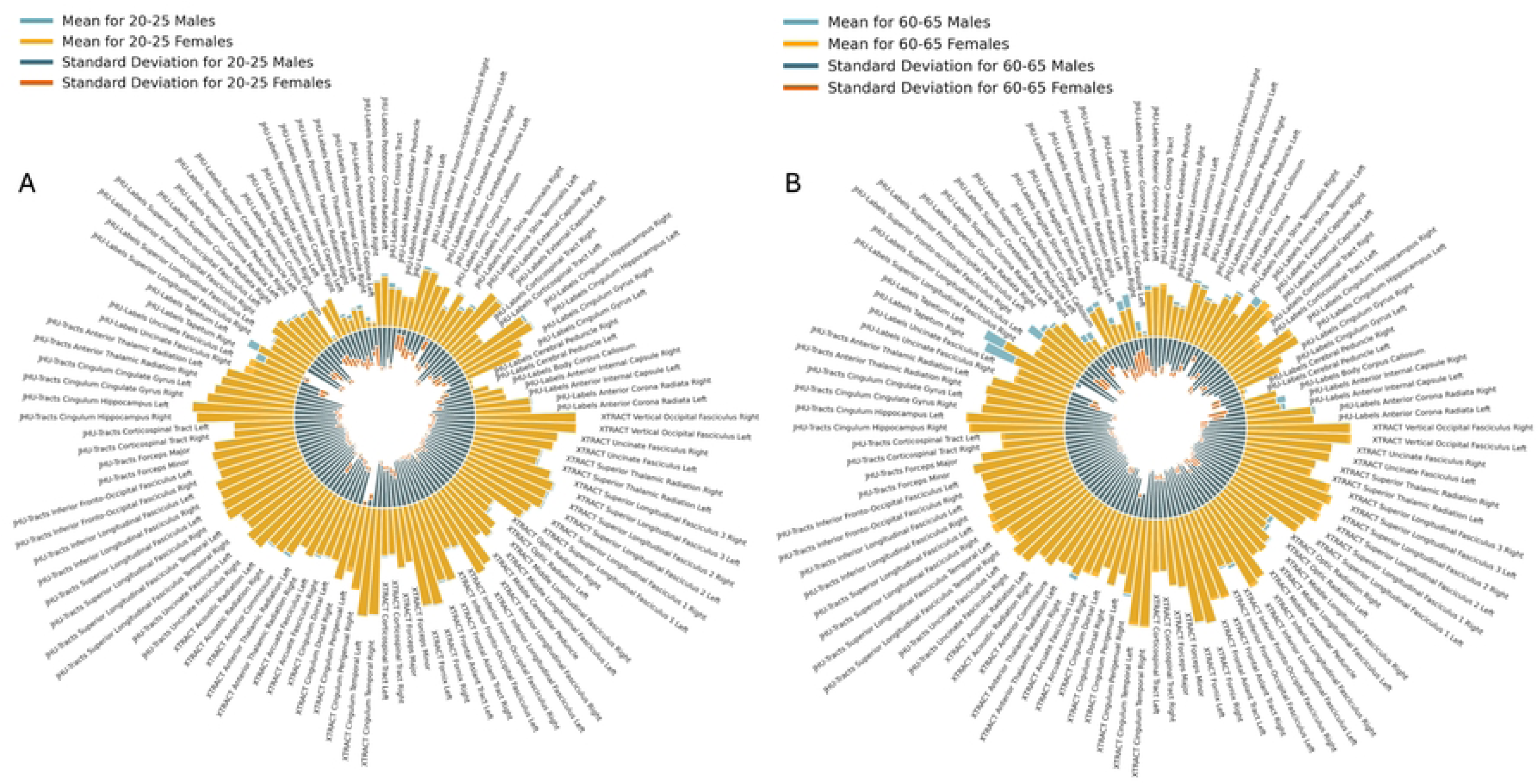
Pair of circular bar plots comparing the mean fractional anisotropy (FA) values and standard deviations across white matter (WM) regions of interest (*ROIs*) for different *Age* and *Sex* groups. In (A), FA metrics for young adults (20-25 years) are displayed while data for older adults are shown in (B). The outer radial plots in both (A) and (B) display mean FA values for males (blue) and females (yellow) across various *ROIs* selected from the JHU-Labels, JHU-Tracts, and XTRACT *Atlases*. The inner radial plot in both (A) and (B) represents standard deviation values for each *ROI* for both males (blue) and females (orange). Generally, males demonstrate higher mean FA values compared to females across most *ROIs*, particularly in the JHU-Labels posterior thalamic radiation (PTR), superior fronto-occipital fasciculus (SFOF), and tapetum of the corpus callosum (CC), and this is more pronounced in older subjects (B). Conversely, standard deviations bars highlight that females show greater variability in FA values across certain *ROIs*, particularly the JHU-Labels anterior internal capsule, PTR, and superior longitudinal fasciculus (SLF). These visualizations indicate significant *Age* and *Sex*-related differences in WM integrity, emphasizing the biological underpinnings and differences in neuroanatomical features related to *Sex* and *Age* in the brain.

## Discussion

The results from this study indicate that significant variability exists between DTI metrics (i.e. FA, MD, AD, RD) and key demographic and scanner-related factors, suggesting that the validity of DTI as a simple biomarker of WM integrity should be evaluated with caution. Furthermore, the study demonstrates that *Age*, *Sex* differences, and technical factors all have relationships with specific WM tracts and anatomical regions which corroborates with existing literature **[20,21]** and thereby emphasizes the need for personalized research approaches and strategic implementation of multi-site and ‘Big Data’ training sets.

Regarding *Age* as a factor, this study confirms that FA decreases with *Age* while AD, MD, and RD increase due to an overall loss in myelin integrity (or quality), a reduction in axonal density, and neurite disorganization **[22,65]**. Cognitive decline and executive function loss is traditionally seen at the fifth decade of life **[56,67]**, but this study highlights that DTI, if controlling for *Age*, *Sex*, and *Vendor* variables, is sensitive to decline that occur starting from the 36-40 year old range **[68,69]**. In younger Age groups (26-30 year old), some slight increases in FA were shown which may be explained by WM maturation **[21]** due to later stages of puberty. Similarly, in adolescence, WM maturation follows a nonlinear trajectory **[21,22]** and this relationship is exhibited inversely in this study. This may be due to healthy aging where WM deterioration follows an almost exponential decay. Results from this study also corroborated the findings that superior regions deteriorate faster with *Age* **[67]**, as manifested in the SLF, SFOF, and SCR. Projection and association type fibers were found to be more susceptible to *Age* degradation, along with the CC. However, other commissural fibers, particularly cingulum-related areas, were more resistant to *Age*-effects. Given these observations in a large healthy population, a valuable quantitative baseline could be created to distinguish between healthy aging and neurodegenerative conditions **[60]**. Future work should target the inclusion of behaviour and cognitive performance **[21]**, emotional regulation, verbal abilities **[22]**, and motor-related tasks to integrate the effects of *Age* on cognitive performance, executive function, and processing speed **[65]**. Longitudinal studies are also warranted as they can help establish clear relationships with critical periods of behavioural decline in aging to further explore the potential of FA as an early indicator of cognitive changes.

This study also highlighted important Sex differences associated with WM integrity. We confirmed that women exhibit lower global FA values than men **[21]**. However, the overall *Age* effect was less generalizable since there were nuances related to the behaviour of specific WM tracts.

Existing literature states that differences exist in the CC and frontal WM **[21]**, with younger males showing higher FA values due to stronger associations with spatial and motor abilities **[22]**. Here we show that relationship exists in younger cohorts, but in older cohorts, females often exhibited higher FA values associated with the CT, the frontal-aslant, and the SLF. As it relates to motor tracts, females possess slower declines in FA suggesting greater resilience to *Age*-related changes and better maintenance of neurite organization **[65]**. Similarly, as women tend to show stronger correlations between emotional and verbal abilities **[22]**, the UF and external capsule, often responsible for emotional regulation, showed much higher FA values in females compared to males. These findings may be linked to biological and hormonal influences, such as estrogen’s neuroprotective effects **[65]** and metabolic changes after menopause **[20]**. However, one of the most interesting findings includes the disparities found between the male and female tapetum of the CC. As females often tend to develop dementia earlier than men **[70]**, it was noted that the tapetum is one of the most significant *ROIs* that demonstrate a large amount of deterioration with *Age*, particularly in females **[71]**. The tapetum may benefit from further exploration as a potential early-stage biomarker related to dementia, Alzheimer’s Disease (AD), and other neurodegenerative diseases. Overall, these findings underscore the importance of considering *Sex* differences in WM study, as they likely contribute to observed variations in cognitive performance, behavior, and vulnerability to neurological conditions.

*Vendor*-related differences in FA measurements were also showcased in this study, emphasizing important considerations in scanner hardware and software configurations in multicenter studies.

Consistent with previous literature **[14,24]**, this study agreed with Siemens scanners generally producing higher FA values compared to GE and Philips. In contrast, Philips machines showed higher FA values in regions such as the forceps minor/major and the ATR, while GE machines tended to report lower FA values in specific tracts like the CT and STR. Additionally, Philips machines were found to exhibit greater variability than any other platform. The *Vendor* findings were further supported by large discrepancies in FA values in regions near the ventricles, which could be attributed to tissue susceptibility artifacts or cerebrospinal fluid (CSF) pulsatile flow (synchronized with the cardiac cycle) potentially inducing additional artifacts **[1]**. Although curation of the dataset included a diffusion direction constraint (> 30), differences in hardware, software, acquisition parameters, motion probing gradients, and resolution size could explain the magnitude of this *Vendor* effect. However, these results emphasize the necessity of considering *Vendor*-specific effects when interpreting FA data, especially in longitudinal, ’Big Data’ or multicenter studies, to ensure meaningful comparisons and reduce potential confounds in clinical and research applications. Moreover, the observed variability underscores the need for further refinement in imaging protocols and the use of standardized quality control (QC) measurements or tools (i.e. phantoms) across sites.

The methodology in the study design of choosing an *Atlas* proved to be another technical factor that played an important role in the overall analysis of the study. The results obtained using the JHU-Labels *Atlas* more consistently corroborated with existing literature compared to the JHU-Tracts and XTRACT *Atlases*. This discrepancy could stem from differences in *ROI* size and volume with larger tracts providing more stable FA estimates than smaller localized regions. The higher sensitivity of the JHU-Labels *Atlas* suggests that future studies aiming to investigate FA-related changes should prioritize larger, well-defined WM tracts over smaller or less anatomically constrained *ROIs*.

Given how technical nuances affected the results in this study, important strategies must be implemented in future work to appropriately use ’Big Data’ and/or multi-site data. Therefore, it is recommended from the results of this study to attempt to limit data collection from only one vendor. Otherwise, it is imperative to implement scanner harmonization techniques **[12,26]** to minimize platform discrepancies. Techniques and tools such as ComBAT **[72]**, protocol standardization, postprocessing correction algorithms, and phantom calibration **[12]** are other alternatives that should be explored to enhance data comparability of diffusion metrics for large-scale imaging research. Future work should also explore scenarios with repeat scans in a multi-site context and investigate the intra-class correlation coefficient (ICC) **[73]**, Dice similarity index **[74]**, and wild bootstrap strategies **[12]** to find more nuanced trends in platform variability to improve future harmonization techniques. It should also be noted that although DTI has become abundant, the technique itself has had major technical developments in recent years, where it is worth investigating other metrics/acquisitions such as neurite orientation dispersion and density imaging (NODDI), q-space imaging, and constrained spherical deconvolution (CSD) analysis that may provide more in-depth findings related to *Age*, *Sex*, *Vendor*, and *Atlas* factors. Overall, these efforts are crucial, particularly in multi-center studies, in providing complementary insights into microstructural alterations beyond FA alone.

## Conclusion

This study comprehensively analyzes the combined contributions of age, sex, ROI, and vendor-related variability to diffusion tensor imaging (DTI) measurements, offering key insights for large-scale, multi-center neuroimaging research. The findings in this study reinforce the impact of biological influences and methodological considerations on white matter microstructure analysis. Healthy aging and sex-specific trajectories, influenced by biological and hormonal factors, underscore the importance of incorporating age and sex as critical variables in neuroimaging. Addressing scanner variability and optimizing atlas selection also remains crucial technical steps for improving the appropriateness and interpretability of diffusion MRI findings in future studies. With DTI being foundational to advancing personalized medicine, the choice of control populations, such as age, sex, and vendor matched cohorts, plays a crucial role in accurately detecting deviations from normative brain patterns **[75]**. This study emphasizes FA as a potential biomarker in ROIs such as the tapetum to enable patient-specific assessments, early disease detection, and targeted therapeutic interventions. Moving forward, integrating standardized imaging protocols within PM frameworks will be essential to ensuring that imaging continues to provide precise, individualized insights into brain structure and function, ultimately contributing to more effective and targeted healthcare solutions.

### Ethics Approval and Consent to Participate

This study was approved by our local research ethics board (Hamilton Integrated Research Ethics Board, HiREB) and the work described was conducted in accordance with the Code of Ethics of the World Medical Association (Declaration of Helsinki) for experiments involving humans.

## Acknowledgments

We thank all open-source data repositories and forums who were instrumental in meeting congruency with our research goals and ensuring proper ethical and diversity requirements were met in this study.

## Credit authorship contribution statement

**Dr. Nicholas Simard:** Conceptualization, Methodology, Software, Validation, Formal analysis, Investigation, Resources, Data Curation, Writing – original draft, Writing – review & editing, and Visualization; **Dr. Michael D Noseworthy:** Conceptualization, Methodology, Validation, Investigation, Resources, Data Curation, Writing – Review & Editing, and Supervision.

## Funding

Funding was provided through a MITACS Accelerate grant to NS and MDN.

## Data Availability Statement

The authors will gladly provide original DICOM data to researchers who send such a request to the communicating author, Dr. M. Noseworthy

## Notes

### Competing Interest Statement

MDN is the CEO and Co-Founder of TBIFinder, Inc.

